# Locomotion-dependent auditory gating to the parietal cortex guides multisensory decisions

**DOI:** 10.1101/2024.02.14.580296

**Authors:** Ilsong Choi, Seung-Hee Lee

## Abstract

Decision-making in mammals fundamentally relies on integrating multiple sensory inputs, with conflicting information resolved flexibly based on a dominant sensory modality. However, the neural mechanisms underlying state-dependent changes in sensory dominance remain poorly understood. Our study demonstrates that locomotion in mice shifts auditory-dominant decisions toward visual dominance during audiovisual conflicts. Using circuit-specific calcium imaging and optogenetic manipulations, we found that weakened visual representation in the posterior parietal cortex (PPC) leads to auditory-dominant decisions in stationary mice. Prolonged locomotion, however, promotes visual dominance by inhibiting auditory cortical neurons projecting to the PPC (AC_PPC_). This shift is mediated by secondary motor cortical neurons projecting to the auditory cortex (M2_AC_), which specifically inhibit AC_PPC_ neurons without affecting auditory cortical projections to the striatum (AC_STR_). Our findings reveal the neural circuit mechanisms underlying auditory gating to the association cortex depending on locomotion states, providing insights into the state-dependent changes in sensory dominance during multisensory decision-making.

## Introduction

Multisensory integration in the brain, which combines neural information from different sensory modalities, is essential for goal-directed perceptual decisions in an ever-changing environment^1^. Animals can make faster^2,3^ and more accurate^4,5^ perceptual decisions when they integrate congruent (coherent) multisensory stimuli. On the other hand, animals frequently prioritize one modality over the other in response to conflicting multisensory stimuli and show sensory dominance in their perceptual decisions^6–13^. For instance, when head-fixed mice faced conflicting audiovisual cues (one instructed ‘Go’ responses while the other instructed ‘No-go’ responses), they often made decisions following auditory cues^12,13^. This sensory dominance in multisensory decisions cannot be fully explained by a simple additive integration of individual stimuli as explained by a classic Bayesian model^14–16^. Rather, it requires under-weighting one modality more than the other before the integration^10,11,17^, potentially via cross-modal inhibition^12,18,19^. Interestingly, the prepotent modality in multisensory decisions not only varies individually but also switches flexibly from trial to trial^10,11,20,21^. Neural underpinnings of such changes in sensory dominance, however, remain largely unexplored.

Perceptual decision-making occurs with sensory representation associated with motor actions across cortical areas^22,23^. The sensory representation is not stable and fluctuates depending on the behavioral states of the animals^24,25^. One key factor affecting the cortical representation of sensory stimuli is the locomotion. Locomotion increases arousal, which amplifies visual gain in the visual cortex (VC)^26–34^ but suppresses auditory representations in the auditory cortex (AC)^35,36^. Furthermore, locomotion generates corollary discharge signals copied from motor commands, which also suppress auditory responses in the AC^37,38^. As a result of these representational changes, mice showed better detection of mild visual cues^27^ and reduced detection of weak auditory cues^35,39^. Despite this, animals still can identify salient auditory cues well and perform auditory-dependent perceptual tasks without disruption even during locomotion^40,41^. These reports suggest that, even during locomotion, animals retain salient auditory information in the brain, enabling them to make appropriate auditory decisions. In a multisensory environment containing both visual and auditory stimuli, it is more challenging to determine how locomotion influences the multisensory representation and decision-making processes. Compared to head-fixed mice, freely roaming mice exhibited no differences in unisensory decisions but only shifted prepotent modality from audition to vision during multisensory decisions^13^. However, it is unclear how locomotion modulates neural circuits representing multisensory stimuli in the brain to alter behavioral decisions in mice.

One of the key brain areas that mediate perceptual decision-making is the posterior parietal cortex (PPC)^42–44^. The PPC encodes various task-relevant activities during visual decision-making tasks. For example, the PPC conveys visual information^45–48^, accumulating sensory evidence^49,50^, history of the task variables^47,51^, animal’s choices^45,47,52–54^, and body movements^55^. Furthermore, inactivation of the PPC resulted in a substantial decline in an animal’s ability to make visual decisions^45,53,56^. Similar to the visual activities, the PPC also showed task-relevant activities during auditory decision-making tasks^47,56–59^. Its inactivation, however, did not impair perceptual decisions based on simple auditory cues, such as feature detection or discrimination in well-trained animals^48,56,60^. Instead, PPC activity was required for more complex auditory-dependent decision-making, which involves higher levels of cognitive processing^58,61,62^. The PPC receives converging inputs from visual, auditory, and somatosensory cortices^63–65^ and responds to multisensory stimuli in humans^66^, primates^19,67^ and rodents^68,69^. However, its role in multisensory decisions is still unclear. PPC inactivation altered multisensory decisions under audiovisual conflicts^12^ (but see also^16^), but it did not affect multisensory decisions in coherent conditions^16,56^. These reports suggest that the PPC in multisensory decisions may depend on the task paradigms or multisensory contexts that the animal encounters.

To understand how the PPC is involved in multisensory decisions in animals making perceptual decisions across auditory, visual, and congruent or incongruent audiovisual stimuli, at different behavioral states, we used *in vivo* calcium imaging and optogenetic manipulation techniques in mice performing an audiovisual discrimination task on the treadmill. We discovered that the dominant modality can be switched from audition to vision during locomotion. The PPC neurons discriminated visual inputs better than auditory inputs, and the visual representation was suppressed when animals made auditory-dominant decisions in audiovisual conflicts. During locomotion, the AC output to the PPC (AC_PPC_) was selectively suppressed, while the AC output to the striatum (AC_STR_) was maintained. This circuit-specific modulation preserved the visual representation in the PPC and allowed animals to make visually dominant multisensory decisions when animals confronted conflicting audiovisual stimuli during locomotion. The locomotion-induced suppression of AC_PPC_ neurons was mediated by top-down inputs from the secondary motor cortex (M2) to the AC. Our data demonstrate a neural mechanism of switching sensory dominance in the face of cross-modal conflicts, which require gating in the auditory-to-parietal circuit according to the animal’s behavioral states.

## Results

### Locomotion enhances visually dominant decisions in mice under audiovisual conflicts

To examine state-dependent multisensory decisions in mice, we trained head-fixed mice to discriminate auditory and visual stimuli and make Go/No-go decisions on a non-motorized treadmill (Fig. 1a,b; 10 kHz vs. 5 kHz pure-tones: auditory Go (A_G_) vs. auditory No-go (A_NG_); vertically vs. horizontally drifting gratings: visual Go (V_G_) vs. visual No-go (V_NG_)). After mice became experts in discriminating both auditory and visual stimuli at similar correct rates in the stationary (S) state (Fig. 1c), we compared their performance in the moving (M) state (Fig. 1d). During the test, either congruent (A_G_V_G_ or A_NG_V_NG_) or incongruent (A_G_V_NG_ or A_NG_V_G_) audiovisual stimuli were randomly presented in a certain ratio (25% or 40%) of trials to examine their licking decisions in this multisensory context (Fig. 1e). During incongruent trials, mice made either auditory-dominant or visual-dominant decisions (Fig. 1e; Methods). Head-fixed mice showed better performance of discriminating congruent audiovisual stimuli compared to auditory-or visual-only stimuli in stationary conditions (Fig. 1f and Supplementary Fig. 1a). In moving sessions, the same mice discriminated the unisensory (auditory or visual) stimuli better, often hit the maximum performances, and therefore showed no further improvement in congruent trials (Fig. 1f and Supplementary Fig. 1a). As reported previously^12^, stationary mice showed auditory-dominant decisions in response to the incongruent audiovisual stimuli (Fig. 1g and Supplementary Fig. 1b; 80.7% of incongruent trials). Interestingly, the moving mice made considerably fewer auditory-dominant decisions (Fig. 1g and Supplementary Fig. 1b; 45.6% of incongruent trials). This enhanced visual dominance (decreased auditory dominance) was not significantly correlated with enhanced visual discrimination (Supplementary Fig. 1c), suggesting that improvements in visual discrimination do not solely account for locomotion-dependent changes in the dominant sensory modality during multisensory decisions.

**Fig. 1.**
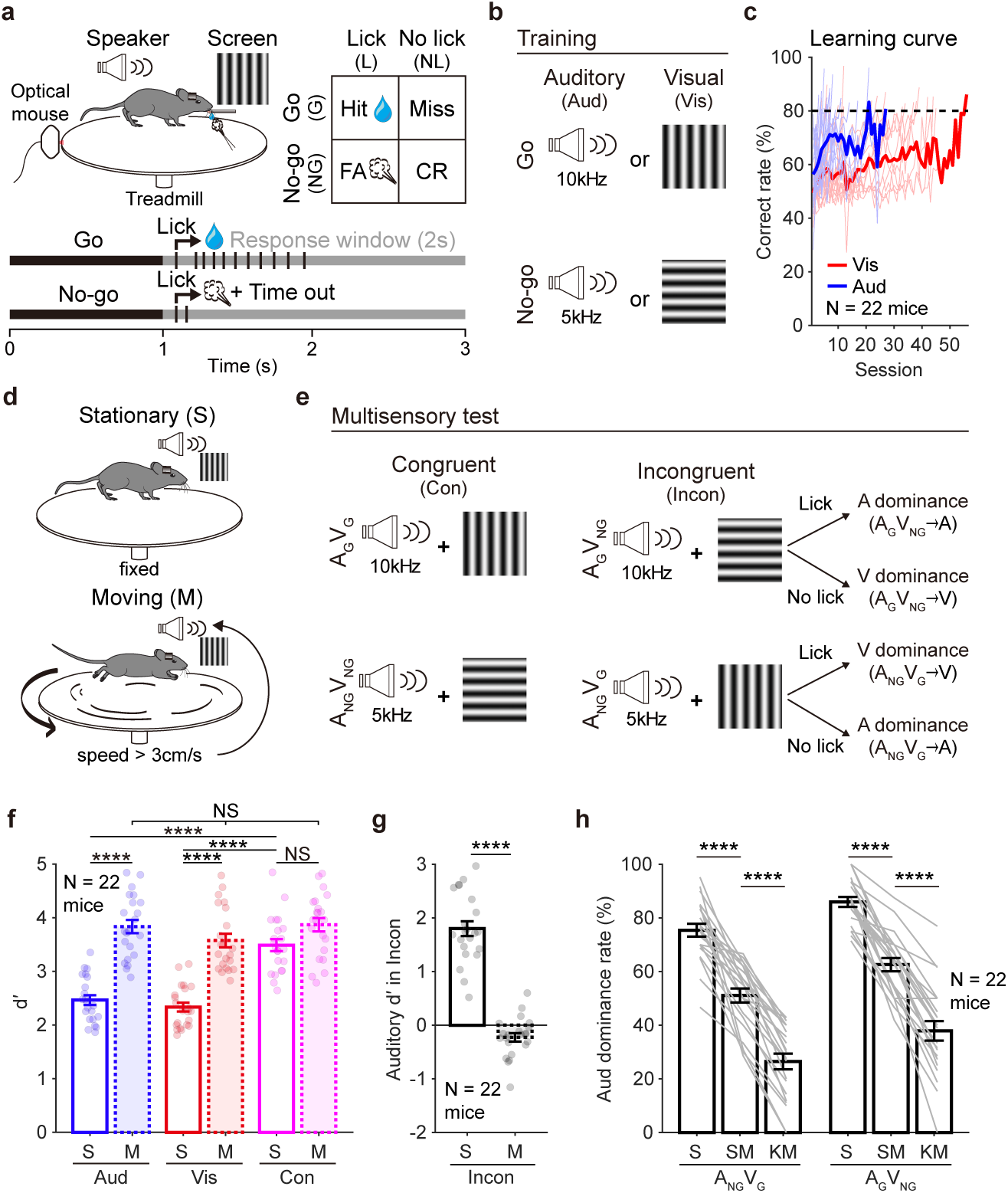
Locomotion shifts auditory-dominant decisions to visually dominant ones in mice under audiovisual conflicts. **a**, Schematic illustration of the audiovisual discrimination task in a head-fixed mouse on a treadmill. G, go; NG, no-go; L, lick; NL, no lick; FA, false alarm; CR, correct rejection. **b**, Stimuli used for behavioral training. Aud, auditory stimuli; Vis, visual stimuli. **c**, Learning curves of mice trained on the task. The black dashed line indicates the criterion for expert performance (80% correct rate). Red, visual discrimination performance; Blue, auditory discrimination performance. **d**, Schematics of stationary (top, S) and moving (bottom, M) conditions. **e**, Audiovisual stimuli used for the multisensory test. Con, congruent stimuli; Incon, incongruent stimuli. Letters ‘A’ and ‘V’ after the arrows indicate auditory and visual dominance, respectively. **f**, Discriminability (d′) for auditory (Aud), visual (Vis), and congruent (Con) Go/No-go stimuli. Each circle represents one mouse. S, stationary; M, moving. **g**, Auditory d′ in mice presented with incongruent (Incon) stimuli. d′ was calculated based on the auditory rules. **h**, Auditory dominance rates in incongruent trials across different locomotion states. Auditory dominance rates reflect the fraction of incongruent trials in which mice followed auditory Go/No-go rules. S, stationary; SM, start-moving; KM, keep-moving. Data are presented as means ± SEM. Thin lines and circles represent individual mouse data. Source data are available in Source Data File. Sample numbers and statistical information are listed in Supplementary Data 1. NS, not significant; \*\*\*\**P* < 0.0001.

We further examined the effects of different conditions of locomotion on switching auditory dominance to visual dominance. First, we examined the effect of locomotion duration, as it has been shown that the locomotion-induced changes in cortical states depend on its duration^36,38,70,71^. We compared mice’s decisions in incongruent trials when the stimuli were presented right after locomotion (start-moving, SM) with those when stimuli were presented after prolonged (> 3s) locomotion (keep-moving, KM). Interestingly, mice showed stronger visual dominance in the keep-moving trials than in the start-moving trials (Fig. 1h and Supplementary Fig. 1b). Once they kept moving at a certain speed (> 3cm/s), the pre-stimulus speed did not further affect their decisions in incongruent trials (Supplementary Fig. 1d). These data indicate that the duration, not the speed, of locomotion is important for switching the dominant modality from audition to vision in mice making decisions under audiovisual conflicts. The unisensory trial types and outcomes of previous trials did not affect the mice’s decision in incongruent trials with or without locomotion (Supplementary Fig. 1e). The level of arousal measured by pupil size was also unrelated to the effect of locomotion on changes in dominant decisions (Supplementary Fig. 1f,g). We also examined whether licking behavior was affected by locomotion (Supplementary Fig. 1h-k). Lick reaction times and post-stimulus lick rates in hit trials were not altered by locomotion (Supplementary Fig. 1i,j), but spontaneous licking before stimulus onset was significantly reduced in the moving state (Supplementary Fig. 1k). However, reduced pre-stimulus licking was not different between auditory-dominant and visual-dominant trials, indicating that pre-stimulus licking did not directly influence the licking decision in multisensory trials (Supplementary Fig. 1k). Therefore, the duration of locomotion prior to stimulus onset, compared to any other internal factors, is the most critical factor determining which modality becomes dominant.

### PPC perturbation enhances auditory dominance and impairs visual discrimination

In our previous study, we found that the feed-forward auditory-to-visual inhibitory circuit in the PPC is critical for mediating auditory dominance in mice performing an audiovisual detection (not discrimination) task in a head-fixed, stationary state^12^. In the present study, we introduced a behavioral paradigm that measured auditory, visual, and audiovisual discrimination in the same mouse. Similarly, we observed auditory dominance during audiovisual conflicts in stationary mice (Fig. 1g,h). Building on these findings, we next examined whether the PPC is required for behavioral decisions during this discrimination task in a stationary state. First, we bilaterally injected muscimol, a GABA_A_ receptor agonist, to suppress PPC activity in task-performing mice (Fig. 2a). PPC inactivation resulted in a severe impairment of visual discrimination but did not affect auditory discrimination in stationary mice (Fig. 2b). This imbalance led to more pronounced auditory-dominant decisions during incongruent trials (Fig. 2b) and reduced multisensory enhancement during congruent trials (Fig. 2c).

**Fig. 2.**
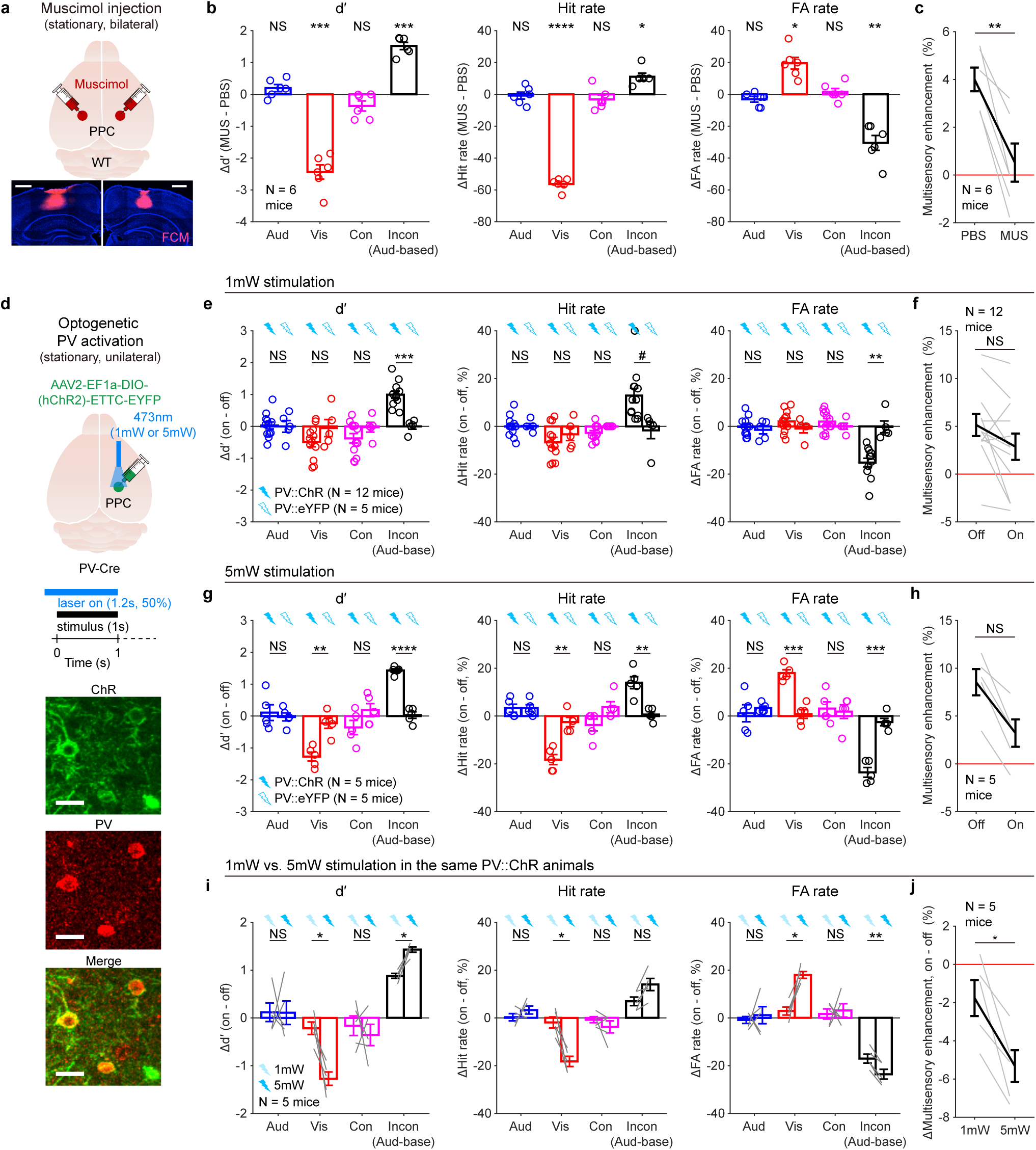
PPC inactivation disrupts visual discrimination and enhances auditory dominance. **a**, Top, schematic illustration of bilateral muscimol injection into the PPC. Bottom, example images showing fluorophore-conjugated muscimol (FCM) injected into the PPC. Scale bars: 500μm. **b**, Changes in d′ (left), hit rates (middle), and FA rates (right) following muscimol injection compared to PBS injection. **c**, Multisensory enhancement in congruent trials after muscimol or PBS injections. **d**, Top, schematic illustration of optogenetic PV^+^ activation in the right PPC. Bottom, fluorescent images showing channelrhodopsin expression in immunostained PV^+^ neurons in the PPC. Scale bars, 20μm. **e**, Changes in d′ (left), hit rates (middle), and FA rates (right) in PV::ChR2 or PV::eYFP mice with 1 mW laser stimulation. **f**, Multisensory enhancement in congruent trials in PV::ChR2 mice with 1 mW laser stimulation. **g** and **h**, same as **e** and **f**, but with 5 mW laser stimulation. **i**, Changes in d′ (left), hit rates (middle), and FA rates (right) with 1 mW or 5 mW laser stimulation in the same PV::ChR2 mice. **j**, ΔMultisensory enhancement (laser on – off; 1 mW vs. 5 mW) in congruent trials. Data are presented as means ± SEM. Circles and gray lines represent individual mouse data. Source data are available in Source Data File. Sample numbers and statistical information are listed in Supplementary Data 1. NS, not significant; *#P* < 0.1, \**P* < 0.05, \*\**P* < 0.01, \*\*\**P* < 0.001, \*\*\*\**P* < 0.0001.

Next, we locally inactivated the PPC in task-performing stationary mice by optogenetically activating parvalbumin-expressing (PV^+^) neurons in the region (Fig. 2d). Mild activation of PV^+^ neurons with a 1 mW laser caused mice to make more auditory-dominant decisions during incongruent trials, while visual discrimination and congruent multisensory enhancement remained intact (Fig. 2e,f). Stronger inactivation of the PPC, achieved using a higher laser power (5 mW), resulted in impairments in visual discrimination and multisensory enhancement similar to those observed under muscimol inactivation conditions (Fig. 2g-j). Interestingly, while robust PPC inactivation impaired visual discrimination, mild activation of PV^+^ inhibitory neurons enhanced auditory dominance without affecting visual discrimination (Fig. 2e,f,i,j), consistent with our previous findings^12^. Collectively, these results demonstrate that PPC inactivation strengthens auditory dominance in multisensory incongruent trials and highlight that PPC activity is essential for visual discrimination but not auditory discrimination.

### Locomotion reduces auditory representation in the PPC

As the PPC activity is important for mice to make perceptual decisions by integrating auditory and visual inputs^12^, we next examined how the PPC excitatory neurons represent task-relevant sensory information in mice performing the audiovisual discrimination task by *in vivo* calcium imaging using a miniature microscope through a prism-combined gradient reflective index (GRIN) lens (Fig. 3a). We first classified neuronal types based on their responses to different sensory stimuli in unisensory trials accompanied by correct decisions (Fig. 3b,c). To avoid any potent influence of the outcome-related activities, we calculated fluorescence changes during the stimulus period before the outcomes (0 ∼ 1 s). In the stationary sessions, many of the PPC excitatory neurons were identified as visual-only (41.7%) or multisensory responsive neurons (36.7%), but relatively small portion of them were auditory-only responsive neurons (10.6%) (Fig. 2c). In the moving sessions, the proportion of visual neurons increased (54.7%), and many neurons lost their auditory responses (Fig. 3c,d and Supplementary Fig. 2a). When we compared the response amplitudes of the PPC neurons, the auditory responses were significantly reduced during locomotion. Still, the visual responses were similar between the stationary and the moving states (Fig. 3d). This finding indicates that locomotion selectively degrades auditory, but not visual responses in the PPC of task-performing mice.

**Fig. 3.**
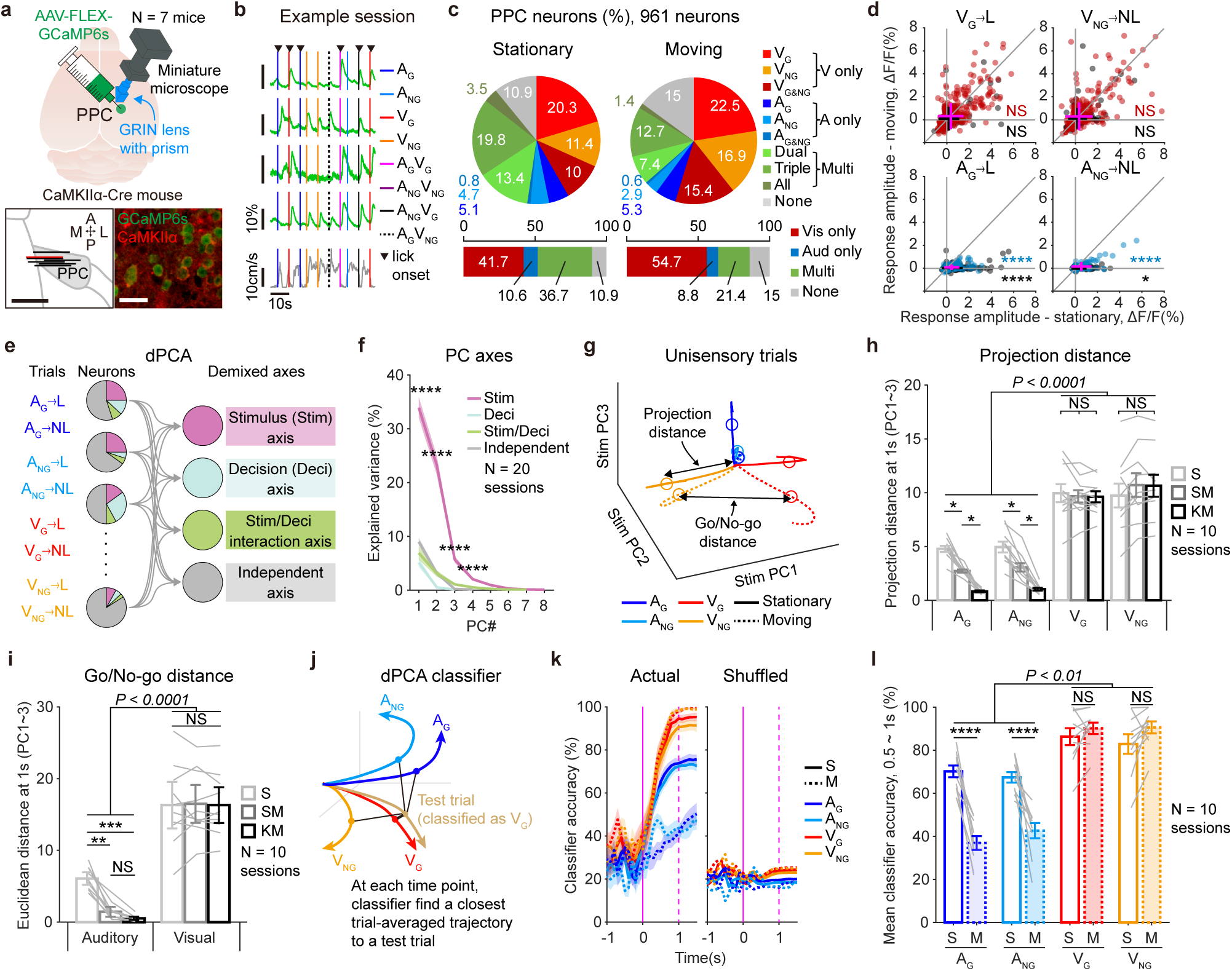
Locomotion suppresses auditory representation while preserving visual representation in the PPC. **a**, Top, schematic of *in vivo* calcium imaging of PPC excitatory neurons in a task-performing mouse. Bottom left, imaging planes across imaged mice (scale bar: 1mm); bottom right, immunohistochemical confirmation of CaMKIIα expression in GCaMP6s-expressing neurons (scale bars: 20 μm). **b**, Calcium activity from four example neurons (green) and locomotion speed (gray) in a moving mouse during the task. **c**, Response types during stationary (left) and moving (right) sessions. Top, percentages of neurons by response type. Bottom, percentages of unisensory and multisensory neurons. **d**, Stimulus-evoked amplitudes of PPC excitatory neurons during correct unisensory trials in stationary vs. moving sessions. Red, visual-selective neurons; blue, auditory-selective neurons; black, multisensory neurons. Magenta (visual and auditory-selective neurons) and black (multisensory neurons) cross-lines indicate means ± SD. **e**, Schematic of dPCA decomposing PPC activity into PC axes representing task-relevant variables. Stim, stimulus; Deci, decision. **f**, Explained variances for the top 20 demixed PCs. Asterisks denote statistical comparisons between stimulus PCs and others. **g**, Stimulus PC trajectories averaged across unisensory correct trials. Circles mark 1 s after stimulus onset. Projection distance, Euclidean distance between points at 0 s and 1 s; Go/No-go distance, Euclidean distance between Go and No-go trajectories at 1 s. **h**, Projection distances of unisensory trajectories. **i**, Go/No-go distances in auditory or visual trials. **j**, Schematic of the dPCA classifier predicting trial types based on Euclidean distances to trial-averaged trajectories. **k**, dPCA classifier accuracy for sorting unisensory trials (left, actual data; right, shuffled trial labels). Solid lines, stationary sessions; dashed lines, moving. For shuffled data, lines represent 95% confidence intervals from 500 repetitions. Colors, unisensory trial types. Magenta, stimulus onset (solid) and offset (dashed). **l**, Mean classifier accuracy during 0.5 ∼ 1 s post-stimulus period. Data are presented as mean ± SEM except for **d**. Gray lines represent individual data. Source data are available in Source Data File. Sample numbers and statistical information are listed in Supplementary Data 1. NS, not significant; \**P* < 0.05, \*\**P* < 0.01, \*\*\**P* < 0.001, \*\*\*\**P* < 0.0001.

During the task, mice often started to lick within the stimulus period, and the stimulus-evoked activity of the PPC neurons might be mixed with the licking-evoked activities during the stimulus presentation period. To decompose the PPC population activity into specific activities representing sensory stimulus, licking decisions, stimulus-decision interactions, or other independent factors, we used demixed principal component analysis (dPCA)^72^. We fed neural data from unisensory trials (i.e., A_G_, A_NG_, V_G_, V_NG_ with licking or no-licking decisions) for the dPCA (Fig. 3e). We confirmed that the decomposed PPC activity in the stimulus PC axes reliably represented each stimulus information regardless of licking actions after the stimuli (Supplementary Fig. 2b). On the other hand, dPCA properly dissociated decision-relevant activity in the PPC neurons in other axes (Supplementary Fig. 2c,d). Interestingly, the stimulus axis explained the PPC population activity much more than any other axes in both stationary and moving conditions (Fig. 3f and Supplementary Fig. 2e), indicating that the PPC neurons represent sensory information the most strongly during the task.

We next examined how the PPC activity represented auditory and visual Go/No-go stimuli in the stimulus subspace by comparing trial-averaged neuronal trajectories of each unisensory stimulus in the top three PCs (Fig. 3g; mean % explanation by top three stimulus PCs: 62.8%). First, we quantitatively measured the magnitude of sensory representations by calculating the projection distance of each trajectory during the stimulus presentation period (for 1 s after stimulus onset) (Fig. 3g; Methods). Auditory trajectories were much shorter than visual trajectories and became even shorter as animals made longer locomotion (Fig. 3h). Next, we calculated the Euclidean distance between a pair of trajectories to quantify similarities in the representation of each stimulus type (Fig. 3i and Supplementary Fig. 2f). Auditory Go and No-go trajectories were close to each other and less distinct during locomotion (Fig. 3i). In contrast, visual Go and No-go trajectories were divergent and distinct from each other (Fig. 3i). The visual trajectories also differed from the auditory trajectories (Supplementary Fig. 2f). Finally, to understand the trial-by-trial reliability of unisensory representation in the PPC, we developed a classifier, which predicted the presented stimulus type in the current trial based on the Euclidean distance between the current-trial trajectory and the trial-averaged trajectory (Fig. 3j). Classification accuracies were significantly higher for the visual trials than the auditory trials (Fig. 3k,l). Moreover, accuracies of auditory classification were significantly lower in the moving sessions than in the stationary sessions (Fig. 3l). Collectively, the PPC neurons processed task-relevant visual information more strongly than the auditory information, showing robust visual but weak auditory Go/No-go discrimination. More importantly, locomotion further dampened the auditory representation in the PPC.

### Multisensory representation in the PPC determines the dominant modality for perceptual decisions under audiovisual conflict

We next examined how the PPC neurons represent multisensory information by projecting their activity into the stimulus subspace during the multisensory trials (Fig. 4a,b; Methods). The neural trajectories representing the congruent multisensory stimuli were very close to the visual trajectories but highly distinct from the auditory trajectories in both stationary and moving conditions (Fig. 4a,b). Euclidean distances, projection distances, and classifier accuracies between the congruent Go and No-go trajectories were not different from those between the visual Go and No-go trajectories regardless of locomotion conditions (Supplementary Fig. 3a-d). These findings suggest that PPC neurons represent congruent multisensory inputs in a similar way that they do visual stimuli.

**Fig. 4.**
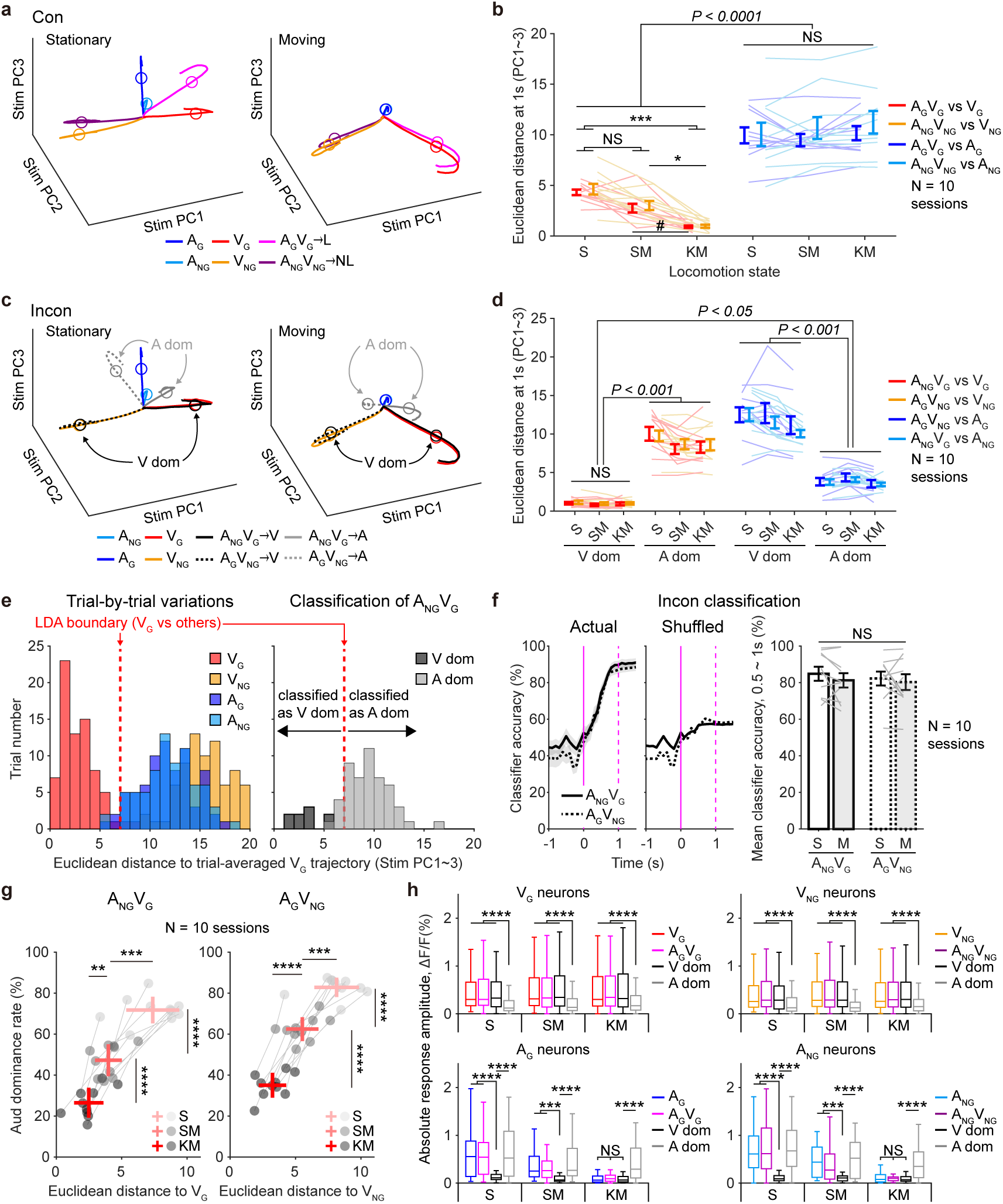
Similarity between visual and multisensory representations in the PPC predicts dominant modality in multisensory decisions. **a**, Trial-averaged calcium trace trajectories for congruent and unisensory trials in the stimulus subspace during stationary (left) and moving (right) sessions. **b**, Euclidean distances between congruent and unisensory trajectories at 1 s after the stimulus onset across locomotion states. **c**, Same as **a**, but for incongruent and unisensory trials. Incongruent trajectories resembled visual trajectories when mice made visual-dominant decisions. Black, visual-dominant trials; Gray, auditory-dominant trials. Solid lines, A_NG_V_G_ trials; dashed lines, A_G_V_NG_ trials. **d**, Same as **b**, but between incongruent and unisensory trajectories. **e**, Left, example session showing Euclidean distances between individual unisensory trajectories and the trial-averaged V_G_ trajectory 1 s after stimulus onset. Linear discriminant analysis (LDA) boundaries were set at distances that maximally separated V_G_ trials from other trials. Right, Euclidean distances between A_NG_V_G_ trajectories and the trial-averaged V_G_ trajectory, using LDA boundaries to classify dominant modalities in A_NG_V_G_ trials. Dark gray, visual-dominant trials; light gray, auditory-dominant trials. **f**, Left, LDA classifier accuracy in predicting dominance in incongruent trials using actual and shuffled trial labels. Solid lines, A_NG_V_G_ trials; Dashed lines, A_G_V_NG_ trials. For shuffled data, lines represent 95% confidence interval upper boundaries from 500 repetitions. Magenta, stimulus onset (solid) and offset (dashed). Right, mean classifier accuracies during 0.5 ∼ 1 s after stimulus onset. **g**, Euclidean distances from incongruent trajectories to visual trajectories and auditory dominance rates at different locomotion states. Left, A_NG_V_G_ trials; right, A_G_V_NG_ trials. Red crosses, means ± SD. **h**, Absolute response amplitudes of visual (top) and auditory (bottom)-responsive neurons during unisensory, congruent, visual-dominant or auditory-dominant incongruent trials at different locomotion states. Data are presented as means ± SEM except for **g**. Circles and connected lines represent individual data. For box plots, central lines indicate medians; boxes represent the 25^th^-75^th^ percentiles; whiskers show maximum/minimum values excluding outliers. Source data are available in Source Data File. Sample numbers and statistical information are listed in Supplementary Data 1. NS, not significant; #*P* < 0.1, \**P* < 0.05, \*\**P* < 0.01, \*\*\**P* < 0.001, \*\*\*\**P* < 0.0001.

By projecting PPC activity trajectories during incongruent multisensory trials, we found that PPC neurons represented the same stimulus condition (A_NG_V_G_ or A_G_V_NG_) differently depending on the impending decisions of mice (Fig. 4c; auditory dominance in grey trajectories vs. visual dominance in black trajectories). We first found that the projection and the Euclidean distances of the trajectories for auditory-dominant incongruent trials were smaller than those for the visual-dominant trials (Supplementary Fig. 3e,f). Furthermore, the trajectories representing the incongruent stimuli were nearly identical to those representing visual stimuli when mice made visually dominant decisions (Fig. 4d and Supplementary Fig. 3g). When mice made auditory-dominant decisions, however, the neural trajectories for the incongruent stimuli diverged from the visual trajectories, becoming closer to but still separate from the auditory trajectories (Fig. 4d and Supplementary Fig. 3g).

We next classified the incongruent trials individually by calculating the Euclidean distance between the single-trial trajectory and the mean trajectories of the four types of multisensory trials and found that the classification accuracy was significantly higher in the stimulus subspace than in other subspaces (Supplementary Fig. 3h). Therefore, the sensory representation in the PPC is the most critical factor determining the multisensory decisions of mice in the audiovisual conflict during the task. We further developed another classifier by setting the boundary of the visual trajectories based on their trial-by-trial variations and classified individual incongruent trajectories as visually dominant ones if they were within the boundary of the visual trajectories (Fig. 4e). Using this, we successfully classified the dominant modality across incongruent trials (Fig. 4f). Collectively, if the PPC neurons dominantly represented visual information under incongruent audiovisual stimuli, mice showed visually dominant decisions. Conversely, if the visually dominant representation weakened in the PPC, mice made auditory-dominant decisions in response to the incongruent stimuli.

To understand if the stimulus representation in the PPC changes with locomotion, we calculated the divergence between the incongruent multisensory trajectory and the visual trajectory at different locomotion states (Fig. 4g). The two trajectories were closer as the animal made longer locomotion, and this was correlated with the decrease in auditory dominance rates in the animal’s multisensory decisions (Fig. 4g). To understand how locomotion altered the trial-averaged neural trajectories in incongruent trials, we further compared the response amplitudes by calculating the absolute values of stimulus-evoked responses in visual-or auditory-responsive neurons across different locomotion and stimulus conditions. Responses of visually responsive neurons were strongly suppressed only in auditory-dominant trials, regardless of locomotion (Fig. 4h top). In contrast, auditory-responsive neurons exhibited stronger responses in auditory-dominant trials compared to visual-dominant trials, while their auditory responses gradually decreased during locomotion (Fig. 4h bottom). Therefore, locomotion increased the likelihood of maintaining strong visual representation while reducing the auditory representation in the PPC and caused mice to make more visually dominant multisensory decisions.

### Locomotion selectively suppresses auditory cortical outputs to the parietal cortex but not to the striatum

The robust reduction of auditory representation in the PPC during locomotion might be due to direct suppression of the auditory neurons in the PPC or indirect suppression of the AC output neurons that project to the PPC (i.e. AC_PPC_ neurons). To examine which case it was, we measured the spontaneous activity changes in the PPC auditory-responsive neurons during locomotion. Neither the locomotion onset nor offset induced significant suppression in their activities (Supplementary Fig. 4a-d). Therefore, the suppression of the auditory responses in the PPC during locomotion is unlikely to result from direct inhibition of the PPC auditory neurons. Instead, it might be due to the reduction of bottom-up auditory inputs to the PPC.

To examine this, we first traced the AC_PPC_ neurons by injecting the retrograde adeno-associated viruses (AAVrg) encoding tdTomato into the PPC (Fig. 5a). We co-traced the AC neurons that project to the posterior striatum (AC_STR_), which are known to play a critical role in the auditory decision-making of mice^73–76^, by injecting the AAVrg encoding eGFP into the striatum (Fig. 5a). AC_PPC_ neurons were mostly found in the anterior and dorsal AC (AuD), whereas AC_STR_ neurons were found across the anterior-to-posterior and dorsal-to-ventral AC (Fig. 5b,c). Only 7% of the traced neurons in the AC were co-labeled, suggesting that the AC_PPC_ and the AC_STR_ neurons are distinct and non-overlapping neuronal populations in the AC (Fig. 5d).

**Fig. 5.**
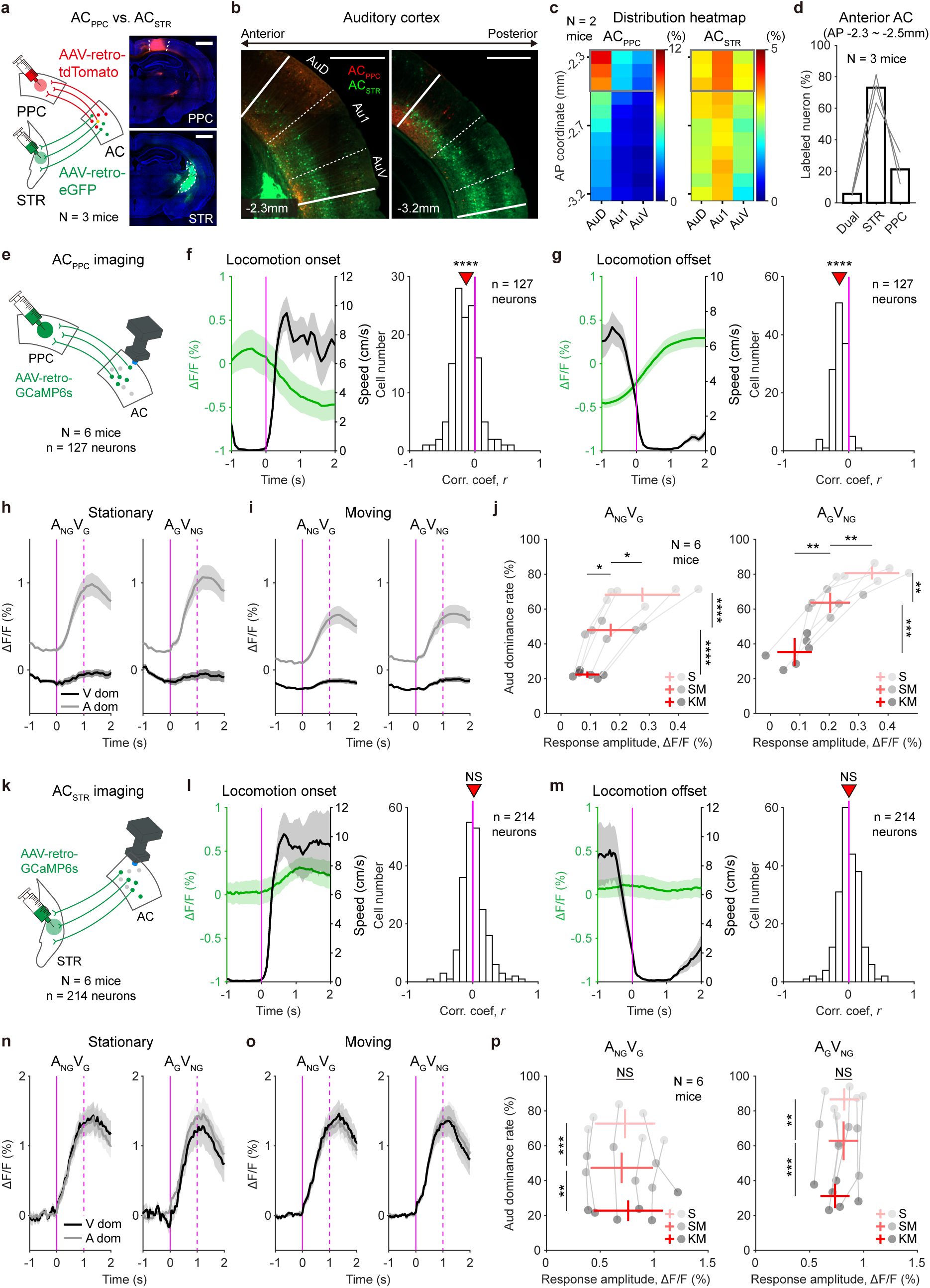
Locomotion preferentially suppresses auditory outputs to the parietal cortex. **a**, Schematic illustration (left) and injection sites (right) of retrograde AAVs for dual retrograde tracing of the AC neurons projecting to the PPC (red, AC_PPC_) and the posterior STR (green, AC_STR_). Scale bars, 1 mm. **b**, Representative images showing AC_PPC_ and AC_STR_ neurons at the anterior (left, -2.3 mm from the bregma) and posterior (right, -3.2 mm from the bregma) AC. Tracing experiments were conducted in 3 mice. White lines, boundaries of the AC; white dashed lines, boundaries between AC subareas. Scale bars, 500 μm. **c**, Heatmaps representing percentages of labeled AC_PPC_ neurons (left) or AC_STR_ neurons (right) in AC subregions at different anterior-posterior coordinates. Gray boxes indicate the anterior AC (-2.3 ∼ -2.5 mm from the bregma). **d**, Percentages of labeled neurons in the AC. Gray lines represent data from individual brain samples. **e**, Schematic illustration of *in vivo* calcium imaging of AC_PPC_ neurons. **f**, Left, mean calcium traces of AC_PPC_ neurons (green) and mean locomotion speeds of mice (black) aligned to locomotion onset. Magenta, locomotion onset. Right, distribution of Pearson correlation coefficients between locomotion speeds and calcium activity of individual neurons at locomotion onset. The red triangle indicates the mean correlation coefficient across all neurons. Magenta, zero line. **g**, Same as **f**, but for locomotion offset. Left, magenta indicates locomotion offset. **h**, Calcium activity of AC_PPC_ neurons during incongruent trials in the stationary state. Black, visual-dominant trials; Gray, auditory-dominant trials. **i**, Same as **h**, but in moving state. **j**, Mean response amplitudes of AC_PPC_ neurons and auditory dominance rates in A_NG_V_G_ (left) and A_G_V_NG_ (right) trials across locomotion states. Circles connected by gray lines represent individual mouse data. Red crosses, means ± SD. **k**-**p**, Same as **e**-**j**, but for AC_STR_ neurons. Data are presented as means ± SEM except for **j**,**p**. Source data are available in Source Data File. Sample numbers and statistical information are listed in Supplementary Data 1. NS, not significant; \**P* < 0.05, \*\**P* < 0.01, \*\*\**P* < 0.001, \*\*\*\**P* < 0.0001.

We next measured the activity of AC_PPC_ and AC_STR_ neurons in mice running on the treadmill (Fig. 5e,k). When animals started to move, the spontaneous activity of AC_PPC_ neurons was significantly reduced, and their calcium activities and locomotion onset speeds were negatively correlated (Fig. 5f). AC_PPC_ neurons were liberated from the inhibition at locomotion offset (Fig. 5g). Unlike the AC_PPC_ neurons, AC_STR_ neurons did not exhibit any activity changes at the onset or offset of locomotion (Fig. 5l,m). These results demonstrate that locomotion suppressed AC_PPC_ neurons directly and selectively without inhibiting AC_STR_ neurons.

We next measured auditory responses in AC neurons across different locomotion states (Supplementary Fig. 5a). Locomotion significantly suppressed the auditory responses of AC_PPC_ neurons but not AC_STR_ neurons (Supplementary Fig. 5b and 5d). When we calculated the stimulus selectivity between the 5kHz and 10kHz tones used for the training, only AC_STR_ neurons showed a significant increase in selectivity during locomotion (Supplementary Fig. 5c and 5e). These data suggest that locomotion selectively suppresses auditory responses in AC_PPC_ neurons while enhancing stimulus selectivity in AC_STR_ neurons, which may be responsible for improved auditory discrimination during locomotion (Fig. 1f).

We also measured visual responses in VC neurons during locomotion (Supplementary Fig. 5f). We identified VC neurons that project to the PPC (VC_PPC_) through retrograde labeling with tdTomato while imaging all VC excitatory neurons in CaMKIIα::GCaMP6f mice using a 2-photon microscope (see Methods). Consistent with previous reports^29,31^, overall VC neurons exhibited increased visual responses and enhanced stimulus selectivity during locomotion (Supplementary Fig. 5g-j). The increased visual responses and stimulus selectivity in VC neurons, including VC_PPC_, may explain the observed improvement in visual discrimination during locomotion. Collectively, our findings indicate that locomotion differentially modulates stimulus-evoked activity in AC and VC neurons, potentially influencing auditory and visual discrimination as well as audiovisual integration.

### Locomotion reduces task-relevant auditory representation in the AC_PPC_ neurons

We next examined the stimulus-evoked activities of AC_PPC_ and AC_STR_ neurons during the task. Both AC_PPC_ and AC_STR_ neurons showed significant auditory responses but no profound visual responses during the task (Supplementary Fig. 6a-c and Supplementary Fig. 7a-c). Prolonged locomotion suppressed the auditory responses of the AC_PPC_ neurons (Supplementary Fig. 6c). In incongruent trials, AC_PPC_ neurons showed strong responses when animals made auditory-dominant decisions but significantly weak responses during visually-dominant decisions (Fig. 5h,i). Locomotion-induced suppression of the AC_PPC_ neurons was correlated with reduced auditory dominance in multisensory decisions of mice (Fig. 5j). However, there was no such suppression in the AC_STR_ neurons during locomotion, and the population of AC_STR_ neurons showed auditory responses at similar levels across the trials independent of locomotion conditions and dominant modalities (Fig. 5n-p and Supplementary Fig. 6c).

We further analyzed AC population activity using the dPCA. As shown in the PPC neurons, the stimulus axis explained the activities of both AC subpopulations more strongly than other task-relevant axes (mean % explanation by top three stimulus PCs: 53.6% for AC_PPC_ neurons; 42.2% for AC_STR_ neurons). During locomotion, the auditory trajectories significantly shrank only in the AC_PPC_ neurons (Supplementary Fig. 6d-f), but not in the AC_STR_ neurons (Supplementary Fig. 7d-f). Furthermore, only AC_PPC_ neurons showed decreased classification accuracies for the auditory stimuli during locomotion (Supplementary Fig. 6g and Supplementary Fig. 7g). During incongruent trials, the response amplitudes and the neural trajectories of the AC_PPC_ neurons were significantly smaller in the visual-dominant trials than in the auditory-dominant trials (Supplementary Fig. 6h-j). In contrast, AC_STR_ neurons showed consistent responses to the incongruent stimuli regardless of dominant modalities (Supplementary Fig. 7h-j). These data indicate that locomotion suppressed the AC_PPC_ neurons more than AC_STR_ neurons, and this selective suppression switched mice to make visually dominant decisions in incongruent trials. In contrast to the vulnerability of the AC_PPC_ neurons, AC_STR_ neurons were unaffected by locomotion, suggesting that the AC_STR_ neurons reliably encode auditory information even during locomotion and are unrelated to multisensory decisions influenced by the locomotion.

### Optogenetic double dissociation reveals distinct roles of the AC subpopulations in perceptual decisions during the audiovisual discrimination task

We wondered whether the two distinct AC outputs, AC_PPC_ and AC_STR_, play distinct functional roles during the audiovisual discrimination task. We first examined whether the activity in the AC_PPC_ neurons was required for auditory dominance in multisensory decisions of mice during incongruent trials. We expressed halorhodopsin (eNpHR3.0) in AC_PPC_ neurons and optogenetically inhibited them in task-performing mice (Fig. 6a). Optogenetic inhibition of the AC_PPC_ neurons did not affect the mice’s decisions in auditory, visual, and congruent audiovisual trials (Fig. 6b). However, it caused a significant reduction in auditory-dominant decisions in stationary mice (Fig. 6b).

**Fig. 6.**
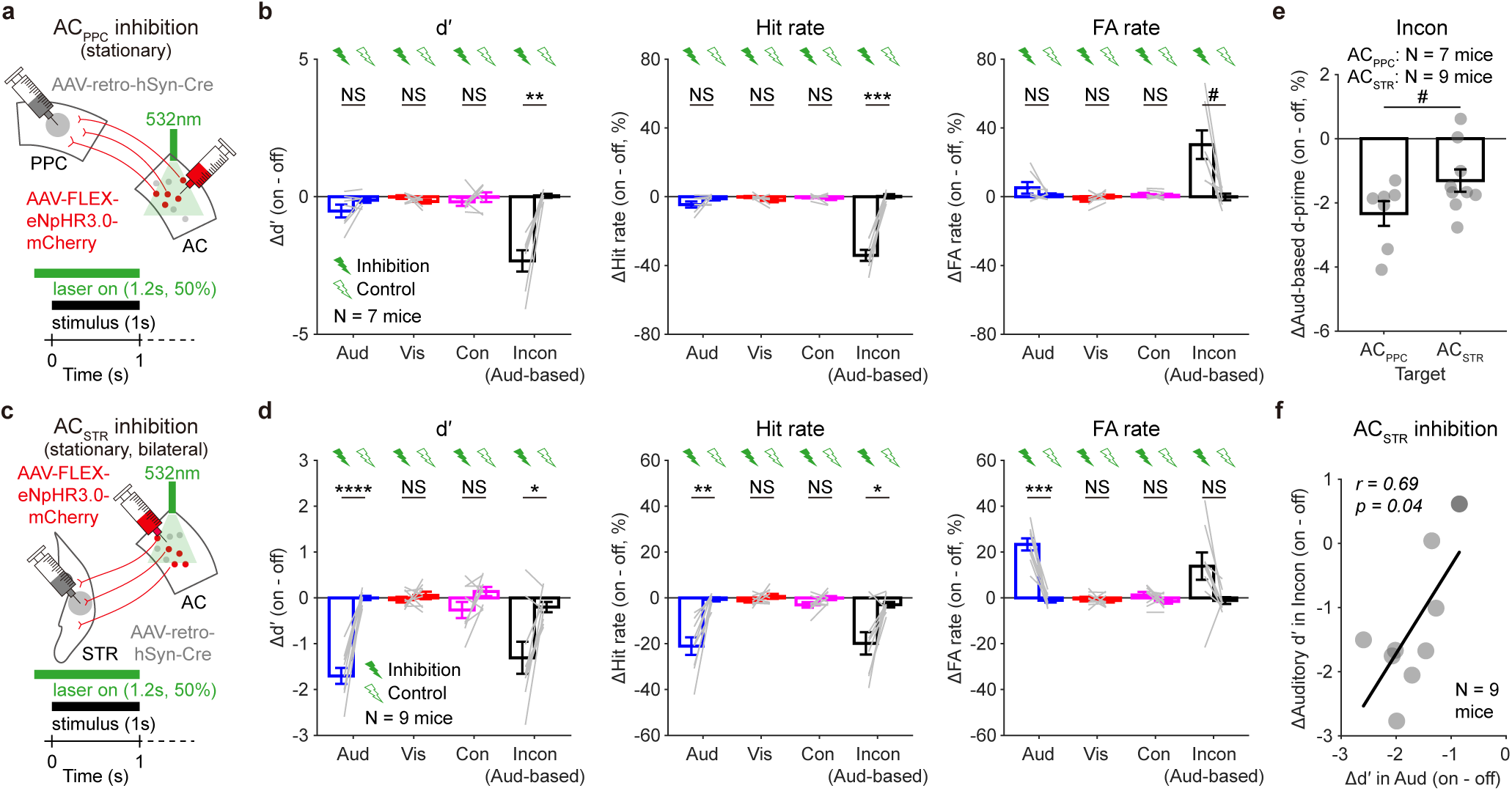
Double dissociation of AC_PPC_ and AC_STR_ neurons in perceptual decision-making. **a**, Schematic illustration of optogenetic inhibition of AC_PPC_ neurons in stationary mice performing an audiovisual discrimination task. **b**, Changes in d′ (left), hit rates (middle), and FA rates (right) following optogenetic inhibition of AC_PPC_ neurons. **c**,**d**, Same as **a**,**b**, but for optogenetic inhibition of AC_STR_ neurons. **e**, Reduction in auditory dominance following inactivation of AC_PPC_ or AC_STR_ neurons. **f**, Correlation between the reductions in auditory discriminability and auditory dominance following inactivation of AC_STR_ neurons. *r*, Pearson correlation coefficient; *p*, significance of the Pearson correlation. Black line, linear regression (slope = 1.37). Bars, means ± SEM. Gray lines and circles represent individual mouse data. Source data are available in Source Data File. Sample numbers and statistical information are listed in Supplementary Data 1. NS, non-significant; *#P* < 0.1, \**P* < 0.05, \*\**P* < 0.01, \*\*\**P* < 0.001, \*\*\*\**P* < 0.0001.

We next examined the role of AC_STR_ neurons during the task by optogenetic inhibition of those neurons (Fig. 6c). Similar to previous studies^73,74^, inhibition of the AC_STR_ neurons significantly hampered auditory discrimination in stationary mice (Fig. 6d). In correlation with a decrease in auditory discrimination, inactivation of AC_STR_ also led to a reduction in auditory-dominant decisions during incongruent trials (Fig. 6e,f). Our findings indicate that the AC-to-STR circuit is important for auditory-dependent perceptual decisions (Fig. 6c,d,f), whereas the AC-to-PPC circuit is more crucial for making auditory-dominant decisions under audiovisual conflicts (Fig. 6a,b,e).

### The auditory cortical inputs inhibit visual representation in the PPC

We next investigated whether AC_PPC_ neurons can suppress visual responses in the PPC during incongruent trials. To address this, we conducted calcium imaging of the PPC excitatory neurons in stationary mice while optogenetically inhibiting AC axons using a miniature fluorescence microscope combined with an optogenetic manipulation system (nVoke; Fig. 7a). Optogenetic inhibition of AC axon terminals in the PPC significantly suppressed auditory responses but did not affect visual responses in the PPC neurons (Fig. 7b). This result indicates that optogenetic inhibition of AC axons in the PPC selectively reduced auditory inputs to the PPC. Next, we compared the response amplitudes of visually responsive PPC neurons between LED-off and LED-on conditions during incongruent trials. As expected, inhibition of AC inputs significantly increased visual responses in the PPC visual neurons (Fig. 7c). Optogenetic inhibition of AC_PPC_ axon terminals also resulted in a significant decrease in auditory-dominant decisions during incongruent trials in stationary mice (Fig. 7d), similar to the effect observed during somatic inhibition of AC_PPC_ neurons (Fig. 6a,b). Collectively, our findings indicate that the AC-to-PPC projection inhibits visual neurons in the PPC, promoting auditory dominance in animals’ decisions during audiovisual conflicts in a stationary state. In contrast, locomotion induces more visually dominant decisions by gating the AC-to-PPC circuit.

**Fig. 7.**
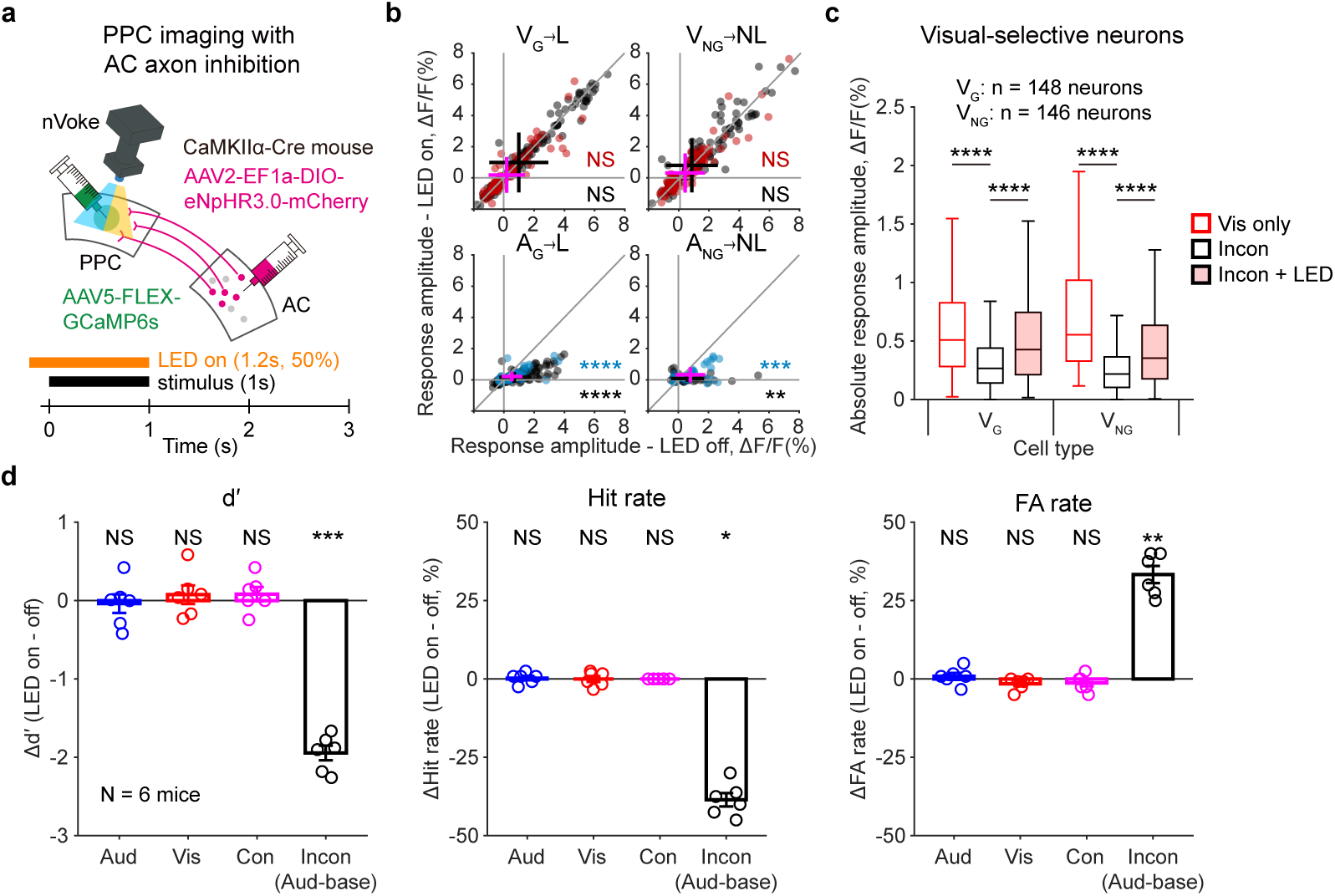
Optogenetic inhibition of AC axon terminals reduces auditory-to-visual suppression in the PPC. **a**, Schematic illustration of *in vivo* calcium imaging with optogenetic inhibition of AC axon terminals in the PPC. **b**, Stimulus-evoked response amplitudes of PPC excitatory neurons in correct unisensory trials across optogenetic LED stimulation states. Red, visual-selective neurons; blue, auditory-selective neurons; gray, multisensory neurons (defined without LED stimulation). Magenta cross-lines, means ± SD for visual-and auditory-selective neurons; black cross-lines, means ± SD for multisensory neurons. **c**, Response amplitudes of visual neurons during visual-only, incongruent, and incongruent trials with LED stimulation in the PPC. **d**, Changes in d′ (left), hit rates (middle), and FA rates (right) following optogenetic inhibition of AC axon terminals in the PPC. Circles represent individual mouse data. Bars, means ± SEM. For box plots, central lines indicate medians; boxes represent the 25^th^-75^th^ percentiles; whiskers show maximum/minimum values excluding outliers. Source data are available in Source Data File. Sample numbers and statistical information are listed in Supplementary Data 1. NS, non-significant; \**P* < 0.05, \*\**P* < 0.01, \*\*\**P* < 0.001, \*\*\*\**P* < 0.0001.

### Motor cortical projection to the AC suppresses AC_PPC_ neurons during locomotion

Previous research reported that the neural projection from the M2 to the AC suppressed auditory responses of AC neurons^37,38^. Our data further showed that locomotion suppressed the AC_PPC_ neurons but not the AC_STR_ neurons. To examine if the M2 mediates this selective suppression of the AC_PPC_ neurons, we first anatomically mapped the spread of fluorescence-labeled M2 axons in the AC (Fig. 8a). Similar to the previous report^37^, M2 axons primarily innervated the superficial or deep layers of the AuD (Fig. 8b). When we compared the M2 axon distribution with the distributions of the AC_PPC_ and the AC_STR_ neurons across AC areas, M2 axons were localized at a similar location where the AC_PPC_ neurons were found (Fig. 8b). On the other hand, majority number of the AC_STR_ neurons did not overlap well with the M2 axons (Fig. 8b). This anatomical feature of the overlap between the M2 axons and the AC_PPC_ neurons in the AuD suggests potential modulation of the AC_PPC_ neurons by the M2 during locomotion.

**Fig. 8.**
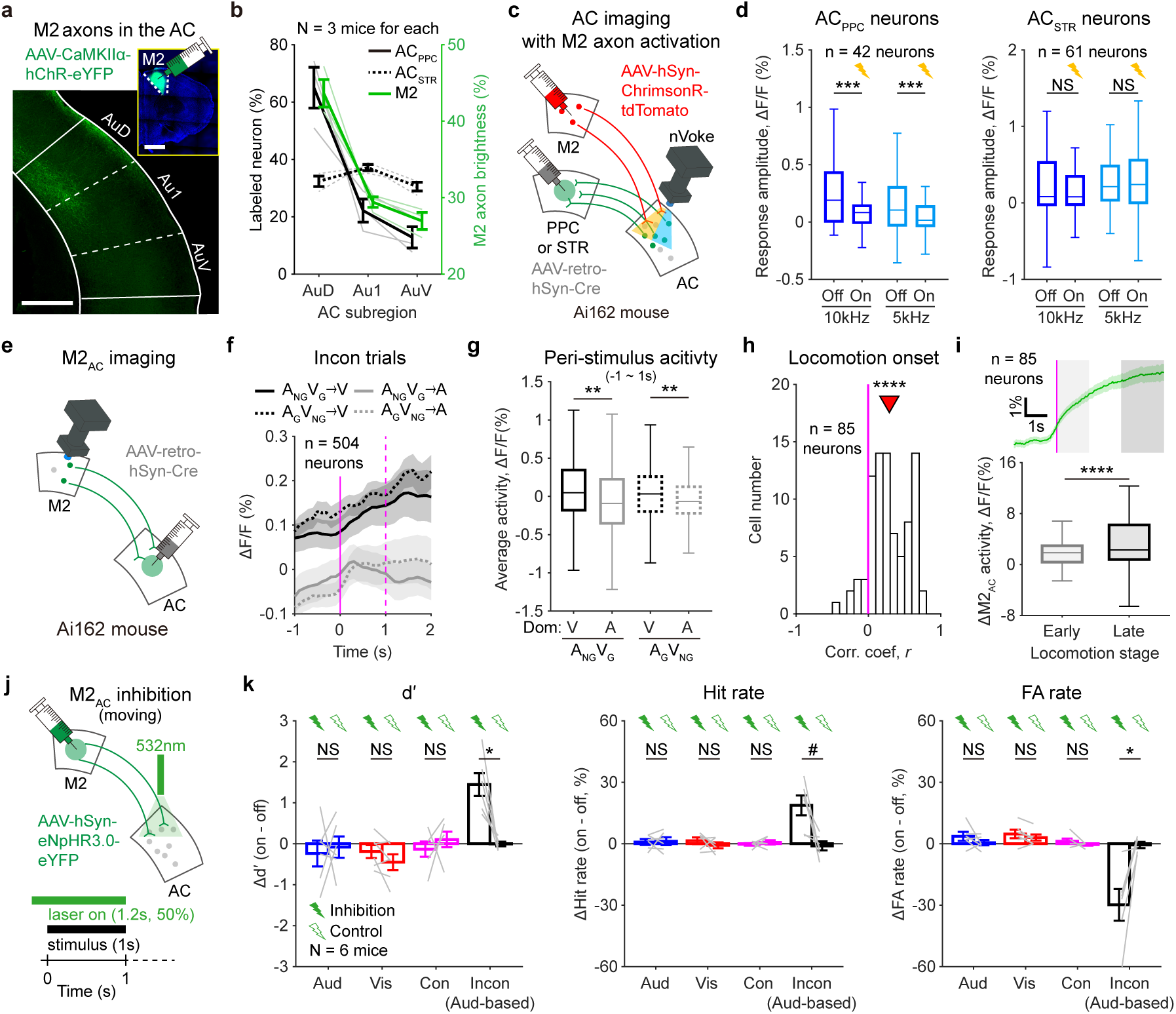
M2-to-AC projection inhibits AC_PPC_ neurons and induces visually dominant decisions during locomotion. **a**, Representative image of virus-expressing M2 axons in subregions of the anterior AC. Scale bar, 500 μm. Inset, an example injection site in M2. Scale bar, 1 mm. **b**, Percentages of labeled neurons (black) and M2 axon intensity (green) in anterior AC subregions. Solid line, AC_PPC_ neurons; dotted line, AC_STR_ neurons. **c**, Schematic illustration of *in vivo* calcium imaging of projection-specific AC neurons combined with optogenetic activation of M2 axons in the AC using the nVoke system. **d**, Response amplitudes of AC_PPC_ neurons (left) and AC_STR_ neurons (right) to 10kHz or 5kHz pure tones. Yellow, optogenetic activation of M2 axons; Off, LED-off; On, LED-on. **e**, Schematic illustration of *in vivo* calcium imaging of M2_AC_ neurons. **f**, Calcium activity of M2_AC_ neurons during incongruent trials. Black, visual-dominant trials; gray, auditory-dominant trials; solid, A_NG_V_G_ trials; dotted, A_G_V_NG_ trials. Magenta, stimulus onset (solid) and offset (dashed). **g**, Peri-stimulus mean activity (-1 to 1 s from onset) of M2_AC_ neurons across incongruent trials. **h**, Distribution of Pearson correlation coefficients between calcium activity and locomotion onset across individual M2_AC_ neurons. Magenta, zero line. **i**, Top, mean activity of M2_AC_ neurons after locomotion onset. Magenta, locomotion onset; lighter gray shade, early locomotion period (0 ∼ 1.5 s after onset); darker gray shade, late locomotion period (> 3 s after onset). Bottom, mean activity of M2_AC_ neurons during early and late locomotion periods. **j**, Schematic illustration of optogenetic inhibition of M2 axons in the AC in task-performing mice during movement. **k**, Changes in d′ (left), hit rates (middle), and FA rates (right) following optogenetic inhibition of M2 axon terminals in the AC. Data are presented as means ± SEM. Thin lines represent individual data. For box plots, central lines indicate medians; boxes represent the 25^th^-75^th^ percentiles; whiskers show maximum/minimum values excluding outliers. Source data are available in Source Data File. Sample numbers and statistical information are listed in Supplementary Data 1. NS, not significant; *#P* < 0.1, \**P* < 0.05, \*\**P* < 0.01, \*\*\**P* < 0.001, \*\*\*\**P* < 0.0001.

We then examined whether this M2 activity suppressed AC_PPC_ neurons directly without affecting AC_STR_ neurons. Using the nVoke system, we activated M2 axon terminals in the AC while imaging calcium activity in AC subpopulations (Fig. 8c). Optogenetic activation of M2 axons significantly dampened the auditory responses of AC_PPC_ neurons (Fig. 8d). In contrast, AC_STR_ neurons were unaffected by M2 axon activation and showed consistent auditory responses during M2 axon activation (Fig. 8d). This result indicates that activation of M2 axons in the AC selectively mediates the suppression of AC_PPC_ neurons.

Next, we investigated whether the activity of M2 neurons that project to the AC correlates with the dominant modality in incongruent trials as well as with locomotion. We first measured the calcium activity of AC-projecting M2 neurons (M2_AC_) in mice performing the task in the stationary condition (Fig. 8e). Interestingly, as shown in the peri-stimulus histogram, M2_AC_ neurons exhibited higher activity in visual-dominant trials compared to auditory-dominant trials (Fig. 8f,g). This result suggests that trial-by-trial variations in the spontaneous activity of M2_AC_ neurons, even before stimulus onset, determine the dominant modality during audiovisual conflicts.

We then measured the calcium activity of M2_AC_ neurons in mice running on the treadmill. M2_AC_ neurons showed robust activation at locomotion onset (Fig. 8h), and the calcium activity of these neurons was significantly higher during the later period of locomotion compared to immediately after locomotion onset (Fig. 8i). This result indicates that M2 neurons copied and delivered the locomotion onset signal to the AC.

Lastly, we optogenetically inhibited the M2 axon terminals in the AC in mice performing the task (Fig. 8j) and found that it did not alter the animal’s decisions in unisensory or congruent trials (Fig. 8k). However, it significantly enhanced auditory dominance in incongruent trials during locomotion (Fig. 8k). These results strongly support the idea that locomotion raises the activity of the M2-to-AC circuit^38^, and we further found that the locomotion-induced activation of this cortical circuit selectively inhibits the AC outputs to the PPC. Consequently, PPC activity is dominated by visual information during locomotion, leading to more visually dominant decisions in mice facing audiovisual conflicts (Fig. 9).

**Fig. 9.**
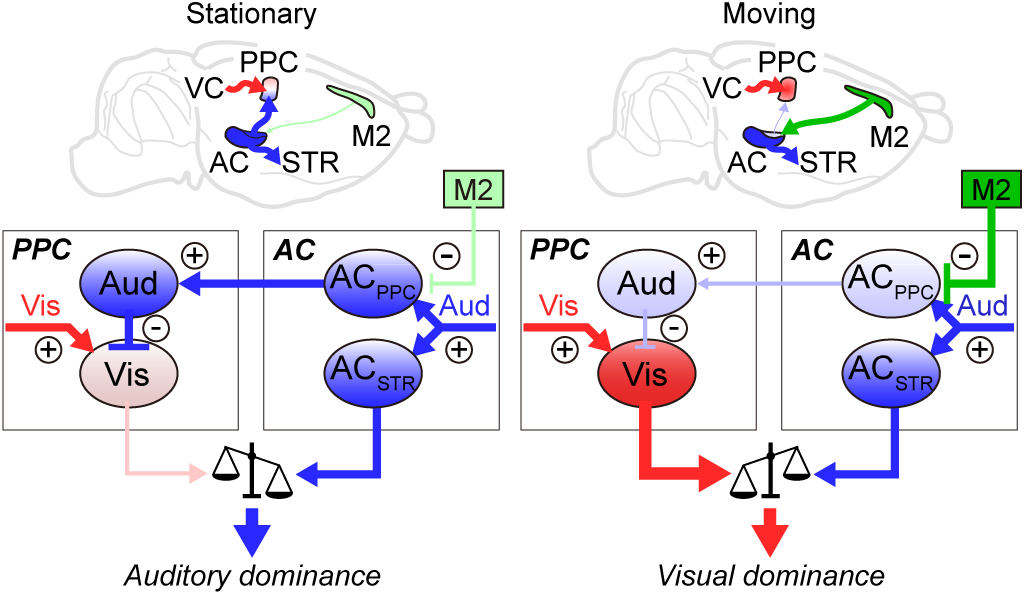
Schematic illustration of sensory dominance switching during locomotion. Left, auditory dominance in stationary mice, driven by auditory-to-visual suppression in the PPC. Right, M2-mediated gating of the AC-to-PPC circuit induces visual dominance during locomotion.

## Discussions

In this study, we identified a neural mechanism underlying behavior state-dependent changes in sensory dominance during multisensory decision-making under audiovisual conflicts in mice. Auditory cortical outputs to the parietal cortex inhibited visual representations in the PPC, leading to auditory-dominant decisions. These auditory outputs were suppressed by locomotion, resulting in more visually dominant decisions. Interestingly, locomotion did not suppress auditory outputs to the STR, allowing animals to maintain precise auditory decisions even during locomotion. Our findings demonstrate that top-down projections from the M2 to the AC selectively gate auditory cortical outputs to the higher association cortex. This gating mechanism specifically regulates how the PPC represents audiovisual conflicts while maintaining accurate perceptual decisions in auditory-only environments in behaving animals.

As all animals naturally locomote while collecting sensory information in the environment to make decisions, understanding the role of locomotion in perceptual decision-making is ethologically critical. More recently, perceptual experiments for head-fixed mice on the treadmill were devised^31^. Using this system, previous studies have examined the effects of locomotion on the simple perceptual detections under the unisensory (either visual or auditory) condition^27,35,39^, but the effect of locomotion on the discrimination of different modality inputs and multisensory decision-making has never been tested. Our study designed a behavioral paradigm to measure perceptual decision-making performances in the unisensory (visual or auditory) and multisensory (congruent or incongruent) conditions at different locomotion states within one session in the same animal. We found that locomotion can improve both visual and auditory perceptual decisions after discrimination of salient stimuli in the same animal. This might be due to the increased stimulus selectivity and sensory responses in the sensory cortices (Supplementary Fig. 5), which can potentially enhance visual and auditory discriminability in animals^34,77^.

We further found that locomotion can switch multisensory decisions from auditory-dominant to visual-dominant ones in mice. The effect of locomotion on multisensory perception and decision has never been examined before. Our findings demonstrate that the dominant modality can be flexibly switched from audition to vision depending on the locomotion states. Sensory dominance in multisensory decisions can be beneficial for animals to make rapid perceptual decisions^78^, which is critical for the survival of animals in a complicated world. Depending on the task design, specific sensory modality information provides more reliable information than other modalities during multisensory integration^78^. A more reliable sensory modality can provide a stronger effect on multisensory decisions in animals performing the task. For example, visual information is more critical for animals to discriminate spatial locations of the multisensory stimuli^9^, while auditory information is more important for temporal discrimination of the stimuli^79^. Our study suggests that the locomotion condition of an animal can flexibly switch the appropriate modality even during the same task. The determination of the dominant modality has been explained by Bayesian models, which explain the optimal strategy of re-weighting different sensory inputs based on their reliability (or variance)^10,11,14^. Our data further suggest that the locomotion generates a self-initiated internal signal that can change the relative weights of the auditory inputs compared to the visual inputs by lowering the reliability of the auditory information in the parietal cortex, where the dominant modality is determined.

Locomotion increases corollary discharge signals in the M2, potentially degrading auditory representations in the AC^38,80^. We further found that locomotion-evoked suppression of auditory responses only happened in the auditory-to-parietal projection (AC_PPC_ neurons) without impairing auditory responses in the auditory-to-striatal projection (AC_STR_ neurons) (Fig. 5 and Supplementary Fig. 6,7). Therefore, corollary discharge signals evoked by locomotion weaken a specific auditory flow from the AC to the higher association cortex. The distinct roles of AC outputs may enable the mammalian brain to maintain auditory processing during locomotion to make auditory-dependent decisions accurately while switching the dominant modality and relying less on auditory stimuli during multisensory decisions under the audiovisual conflicts.

Interestingly, we also found that M2_AC_ neurons exhibited spontaneous activity fluctuations that correlated with visual dominance rates even in stationary mice (Fig. 8e-g). This activity fluctuation may reflect internal state changes that are independent of actual motor output before locomotion or other unmeasured motor behaviors in head-fixed mice. Future studies are required to determine whether M2 activation specifically inhibits the dorsal AC, where AC_PPC_ neurons are located, by forming specific top-down inhibitory synaptic connections, rather than affecting other cortical areas or neurons.

In contrast to auditory representation in the PPC, we found that locomotion did not significantly weaken or strengthen visual representation in the PPC (Fig. 3). However, some individual neurons reshaped their visual responsiveness (either increased or decreased, Supplementary Fig. 2a) during locomotion. Furthermore, most VC neurons showed enhanced visual responses and stimulus selectivity during locomotion (Supplementary Fig. 5f-j), possibly due to cholinergic activation in the VC^29^. This improved visual processing may lead to better visual discrimination during locomotion. However, the increase in visual dominance during multisensory trials was independent of enhanced visual discrimination during locomotion (Supplementary Fig. 1c), suggesting that these two modulations occur in parallel via different pathways.

Consistent with previous research^48^, we found the PPC excitatory neurons represent visual information more reliably than auditory information during the task. Although we observed more visually responsive neurons than auditory-responsive neurons in the PPC, our previous study showed balanced auditory and visual responses in the PPC (see Fig. 7 of reference^12^). This difference was mainly due to increased visual responses in PPC neurons compared to the previous study. The increased portion of visual neurons in this study might be attributed to differences in stimulus types (complex drifting grating vs. optogenetic cortical stimulation and flash), task-learning states (expert vs. naïve mice), and anesthesia states (awake and task-engaged vs. anesthetized) between the current and previous studies. Despite this discrepancy, both studies agree that strong auditory inputs to the PPC can disrupt its visual representation, thereby facilitating auditory-dominant multisensory decisions in head-fixed, stationary mice.

Interestingly, PPC excitatory neurons discriminated auditory Go/No-go information during auditory-based decisions less effectively than visual information, suggesting that the PPC itself is not involved in auditory decision-making. Supporting this, when we inactivated the PPC either through muscimol injection or strong optogenetic activation of PV^+^ inhibitory interneurons, we observed impairments in visual discrimination but not auditory discrimination behaviors. Therefore, the PPC is more critical for visual decision-making in mice. Under multisensory conditions, the PPC integrates auditory inputs that can negatively impact visual weighting, and such integration is necessary for auditory-dominant multisensory decisions in incongruent trials. We also found that the AC output to the STR, which is important for auditory decision-making, exhibited stable auditory representation even during visually dominant decisions (Fig. 5 and Supplementary Fig. 7). Future studies are required to fully understand whether the strong visual representation in the PPC is also required to suppress downstream areas of AC_STR_ neurons to shift auditory-dominant decisions to visually dominant ones. This unknown circuit might be responsible for the visual dominance observed when the PPC was inactivated, as discussed in our previous study^12^.

Our data also indicate that the suppression of visual responses in the PPC only occurs in incongruent trials (not congruent trials) when animals make auditory-dominant decisions (Fig. 4h). One potential explanation for this congruency-dependent cross-modal suppression would be an antagonistic selectivity between Go and No-go stimuli in auditory-to-visual inhibitory circuits. Supporting this idea, opponent inhibition between choice-selective neurons was recently found in the PPC^81^. Our previous report showed that PV^+^ neurons in the PPC strongly received AC inputs and mediated auditory-dominant behaviors^12^. Future studies are required to examine whether the selectivity of inhibitory neurons emerges with learning in these inhibitory neurons in the PPC. Similar to the role of the mouse PPC exerting cross-modal inhibition, the human prefrontal cotex^82^ and the superior temporal sulcus^17,83^ also showed multisensory depression in resolution of conflicting audiovisual speech stimuli. Therefore, multisensory depression might happen across association areas when animals face multisensory conflicts, and our study demonstrates how locomotion states flexibly adjust such inhibition during multisensory conflicts by gating bottom-up inputs.

## Methods

### Animals and surgery

#### Animals

All experimental procedures were approved by the KAIST Institutional Animal Care and Use Committee (IACUC-18-100, IACUC-18-235). Both male and female mice (P45-P150) were used in this study. Animals were maintained *ad libitum* under light (8 am-8 pm) and dark cycle (8 pm-8 am) conditions and housed under single animal per cage conditions from the start until the last day of the experiment (temperature: 20 ∼ 22°C; humidity: 30 to 50%). We used C57BL/6J wild-type (WT) mice, B6;129P2-Pvalb^tm1(cre)Arbr^/J mice (PV-IRES-Cre, stock no. 008069, Jackson Laboratory), B6.Cg-Tg(Camk2a-cre)T29-1Stl/J mice (CaMKIIα-Cre, stock no. 005359, Jackson Laboratory), Ai148 mice (stock no. 030328, Jackson Laboratory) crossed with the CaMKIIα-Cre mice, and Ai162 mice (stock no. 031562, Jackson Laboratory).

#### Common procedure of animal surgery

Adult mice (P45-P80) were anesthetized with isoflurane (1.5%–2% in oxygen) and fixed their head on the stereotaxic apparatus. The body temperature was monitored and maintained at 37°C using a heating pad connected to a body temperature sensor. The scalp was alternately sterilized three times with 70% ethanol and povidone-iodine solution. After the sterilization of the scalp, surgical scissors was used to cut the scalp according to the midline (for virus injection) or remove a portion of the scalp (for the implantation of head plates, GRIN lens, and optic fiber). Periodsteum above the skull was removed, and povidone-iodine solution was used to sanitize the exposed skull and incision. Lastly, we cleaned the exposed skull with sterilized phosphate-buffered saline (PBS).

#### Virus injection for retrograde tracing

For the retrograde tracing of the AC neurons that project to the PPC and the STR, we injected 0.5 μl of retrograde AAV expressing tdTomato (AAVrg-CAG-tdTomato, Addgene) and 0.5 μl of retrograde AAV expressing GFP (AAVrg-CAG-GFP, Addgene) into the target areas respectively. After 2∼3 weeks from the injections, we perfused mice to collect the brain samples for histology.

#### Head plate implantation surgery for mouse head-fixation

To implant a stainless steel head-plate on the mouse head, three miniature stainless steel screws (J.I. Morris) were implanted into the exposed skull using cyanoacrylate glue (Loctite super glue 401, Henkel). A customized stainless steel head-plate was attached to the skull with the cyanoacrylate glue, and the between the skull and head-plate was filled by dental acrylic (Ortho-Jet, Lang Dental).

#### Surgical procedures for in vivo calcium imaging with miniature microscope

To express GCaMP6s in the target neurons, we made a small craniotomy (diameter ∼0.5 mm) above the targeted location. To conduct *in vivo* calcium imaging at the PPC, we injected ∼0.5 μl of Cre-recombinase dependent adeno-associated virus (AAV) expressing GCaMP6s (AAV5-Syn-Flex-GCaMP6s-WPRE-SV40, UPENN) into a site that is 0.2 mm away from the right PPC in posterior direction (posterior 2.15 mm, lateral 1.6 mm from bregma; depth 0.35 mm) of CaMKIIα-Cre mice using a thin pulled glass pipette (20∼25 μm of tip diameter, item no. 504949, WPI) installed in a Nanoliter 2010 injector (WPI) at a speed of 24 nl/min. For *in vivo* calcium imaging of AC_PPC_ neurons, we injected ∼0.5 μl of retrograde AAV expressing GCaMP6s (AAVrg-hSyn1-GCaMP6s-P2A-nls-dTomato, Addgene) into the right PPC (posterior 1.95 mm, lateral 1.6 mm from bregma; depth 0.35 mm) of the WT mice. For *in vivo* calcium imaging of AC_STR_ neurons, we injected ∼0.5 μl of retrograde AAV expressing GCaMP6s (AAVrg-hSyn1-GCaMP6s-P2A-nls-dTomato, Addgene) into the right STR (posterior 1.5mm, lateral 3.45mm from bregma; depth 2.8mm) of the WT mice. For *in vivo* calcium imaging of AC-projecting M2 neurons, we injected ∼0.5 μl of retrograde AAV expressing Cre-recombinase (AAVrg-hSyn-Cre, Addgene) into the right AC (posterior 2.5mm, lateral 4.2mm from bregma; depth 0.8mm) of the Ai162 mice. For *in vivo* calcium imaging of the PPC neurons with optogenetic inhibition of AC axons with nVoke system (Inscopix), we injected ∼0.5 μl of AAV5-Syn-Flex-GCaMP6s-WPRE-SV40 into the right PPC (posterior 2.15 mm, lateral 1.6 mm from bregma; depth 0.35 mm) and injected Cre-recombinase-dependent AAV expressing eNpHR3.0-mCherry (AAV2-EF1α-DIO-eNpHR3.0-mCherry, UNC Vector Core) into the right AC. For *in vivo* calcium imaging of AC neruons with optogenetic of M2 axons with nVoke system, we injected ∼0.5 μl of retrograde AAV expressing Cre recombinase into the right PPC or STR and ∼0.5 μl of AAV-hSyn-ChrimsonR-tdTomato (Addgene) into the right M2 at two anterior-posterior coordinates (anterior 1.0 & 1.7 mm, lateral 0.75 mm from bregma; depth -0.6 mm) of Ai162 mice. After the injection, the incision was sutured with a stitching fiber and disinfected with a povidone-iodine solution.

After 10-14 days from the virus injection, gradient reflective index (GRIN) lens implantation surgery was conducted as previously demonstrated^84^. Mice received subcutaneous injections of carprofen (5 mg/Kg) and dexamethasone (5 mg/Kg) 1 hour before the surgery. After anesthetization of a mouse, a 1 mm^2^ of square-shaped craniotomy was made on the skull above the area of implantation. A prism-attached GRIN lens (1050-004601 or 1050-004606, Inscopix) was inserted 0.2mm away from the imaging area, where we made a small linear incision of the cortical surface using a surgical knife (10055-12, Fine Science Tools) coupled to the stereotaxic apparatus. The GRIN lens was gently inserted (-0.1 mm/20 s, depth 0.8 mm for the PPC; 1.4 mm for the AC; 1.1 mm for the M2) in different directions of the image plane based on the target areas (PPC and AC: along the medial-lateral axis; M2: along the anterior-posterior axis). The exposed brain tissue between the GRIN lens and skull was covered with Kwik-Sil (WPI), and the inserted GRIN lens was secured by applying Super-Bond (Super-Bond C&B Kit, Sun Medical Co.) and the cyanoacrylate glue over the cured Kwik-Sil layer and connected skull. Mice were then implanted with the head plates. Mice were subcutaneously injected with carprofen and dexamethasone for three days following surgery. The field of view of each mouse was examined on a regular basis, and if we detected obvious neuronal activity, we fastened a baseplate (1050-004638, Inscopix) atop the GRIN lens using dental acrylic. To protect the lens from dirt, the baseplate was covered with a baseplate cover (1050-004639, Inscopix).

#### Surgical procedures for in vivo calcium imaging with 2-photon microscope

We injected 0.5 μl of retrograde AAV expressing tdTomato (AAVrg-CAG-tdTomato, Addgene) into the right PPC of cross-line mice (Ai148 x CaMKIIα-Cre). 10∼14 days later, craniotomy and head-plate implantation surgery were conducted as previously demonstrated^85^. Mice received subcutaneous injections of carprofen (5 mg/Kg) and dexamethasone (5 mg/Kg) 1 hour before the surgery. After the anesthetization of a mouse, a round craniotomy with a 3 mm diameter was made on the right posterior cortical area. A 3-layered round cover glass, which was made of 2 pieces of 3-mm-diameter cover glass (W4 64-0720, Warner Instruments) and 1 piece of 5-mm-diameter cover glass (W4 64-0700, Warner Instruments) stacked by an optical adhesive (no. 71, Norland products), was inserted into the craniotomy, and the edge of the glass window was glued with VetBond (B00016067, 3M) and Super-Bond. Mice were then implanted with the head plates. Mice were subcutaneously injected with carprofen and dexamethasone for three days following the surgery.

#### Surgical procedure for optogenetic manipulation during behavior task

For optogenetic activation of PV^+^ neurons in the right PPC, we injected 0.5 μl of Cre-recombinase-dependent AAV expressing ChR2-eYFP (AAV2-EF1a-DIO-hChR2-ETTC-eYFP, UNC Vector Core) or eYFP (AAV2-EF1a-DIO-eYFP, UNC Vector Core) into the PPC. For optogenetic inhibition of projection-specific AC neurons, we bilaterally injected retrograde AAV expressing Cre-recombinase (AAVrg-hSyn-Cre-WPRE-hGH, Addgene) into the right PPC or bilateral STR (0.5 μl per each site) and 0.5 μl of Cre-recombinase-dependent AAV expressing eNpHR3.0-mCherry (AAV2-EF1a-DIO-eNpHR3.0-mCherry, UNC Vector Core) into the AC. For optogenetic inhibition of the M2-to-AC circuit, we injected the excitatory-neuron-specific AAV expressing ChR2-eYFP (AAV2-CAMKIIa-ChR2-EYFP, UNC Vector Core) or the AAV expressing eNpHR3.0-eYFP in neurons (AAV2-hSyn-eNpHR3.0-eYFP, UNC Vector Core) into the right M2 at two anterior-posterior coordinates of WT mice (0.5 μl per each coordinate). After the injection, mice were further implanted with the head plates. Mice had 14 days of viral expression before starting water restriction for the behavioral experiment. When mice were trained and ready for the manipulation experiment (see Methods; **Behaviors**), we provided mice with sufficient water for 2 days and implanted optic fiber on target sites for the laser light delivery. Mice were anesthetized with 1.5-2 % isoflurane, and a 0.5 mm-diameter craniotomy was made on the right AC. An optic fiber with a 200 µm diameter was implanted into the right AC (0.6mm depth) and glued with Kwik-Sil and cyanoacrylate glue. For the AC_STR_ inhibition (Fig. 6c), we implanted optic fibers into the AC bilaterally. We recovered mice for three days after the surgery and restarted the water restriction and behavioral training.

### Behaviors

#### Hardware setups for behavior experiments

Visual stimuli were generated with a custom MATLAB code. A gamma-corrected LCD monitor (12 cm x 9 cm; minimal luminance: 0 lux; maximal luminance: 120 lux; refresh rate: 60Hz) was placed 10 cm away from the left eye of mice at a 45° angle from the rostral-caudal axis. For the binocular visual stimulation task, we put another monitor in front of the right eye of mice as we put on the left side. Full-field drifting gratings (100% contrast, 2 Hz temporal resolution, 0.04 cycles/° spatial resolution, 60 Hz refresh rate) were used for the visual discrimination task in two directions (rightward for Go; upward for No-go). Auditory stimuli were generated with a custom MATLAB code (sampling rate 119,000 Hz, 16bit). A speaker (#60108, Avisoft) was placed 10cm away from the left ear of the mice. Pure tones with different frequencies (10 kHz for Go, 5 kHz for No-go in the main task) were used for the auditory discrimination task. All sound stimuli were calibrated to have a 76 dB sound pressure level (SPL) at the left ear of mice.

Licking behavior of task-performing mice was detected by a custom-made lickometer. We delivered water reward and air punishment via solenoid valves (EV-2-24, Clippard). All the hardware components were operated and controlled by Presentation (Neurobehavioral Systems) with USB data acquisition devices (USB-201, Measurement Computing). An optical mouse (G100s, Logitech) was placed beside a spinning disk of the treadmill. The rotation of the disk during locomotion was converted to digitalized signals by the optical mouse, and real-time locomotion speed was calculated by a customized LabView code (sampling rate: 40 Hz). If the speed exceeded the locomotion threshold (3 cm/s), a 5V-digital signal was sent to the Presentation software in real time through a USB data acquisition device (USB-6001, National Instruments) to present sensory stimuli during locomotion.

### Behavioral training and multisensory discrimination experiments

Mice underwent water restriction (∼1 ml of water per day) throughout the behavioral experiment. The weight of mice under the water restriction was monitored throughout the experiments and maintained at more than 75% of the initial body weight before the water restriction. We calculated the correct rate, hit rate, false alarm (FA) rate, and auditory dominance rate as follows:

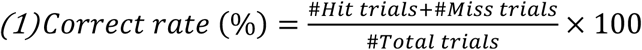

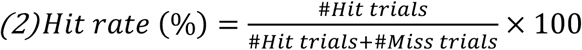

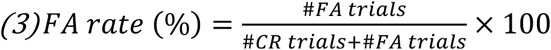

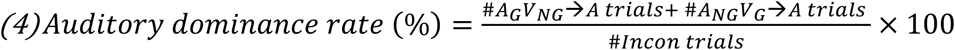

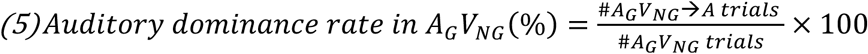

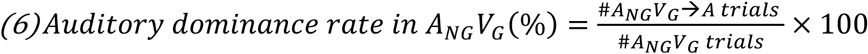

We used adjusted hit rate and FA rate to calculate d′ from behavior data to avoid infinite values due to the 100% hit rate or 0% FA rate in some sessions.

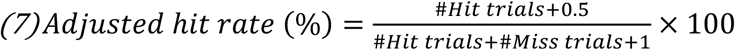

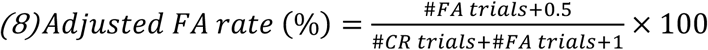

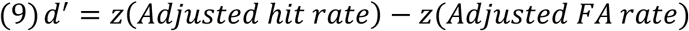

All mice were trained and tested through the following five steps: 1) reward conditioning, 2) stimulus conditioning, 3) audiovisual discrimination training, 4) treadmill habituation; 5) audiovisual integration test

1. Reward conditioning: water-restricted mice received 0.8 μl of water reward whenever they licked the water port. If mice did not show any spontaneous licking behavior within the first 3 minutes after the beginning of a session, we manually delivered water through the water port until mice started to lick the water port. If mice got more than 0.8 ml of total water reward (> 1000 licks) within the first 20 minutes without manual water delivery, mice were moved to the second step in the next session.
2. Stimulus conditioning: auditory go stimulus, visual go stimulus, and no stimulation trials were pseudo-randomly presented in the proportion of 1:1:2. If mice licked the water port during 2 s of response window after the stimulus offset, 1 μl of water reward and 2 s of reward consumption time were presented (hit). If mice did not lick the water port until the offset of the response window (miss), 1 μl of water reward was delivered at the end of the response window to encourage learning. If mice licked the water port during the no stimulation trial (FA), 6 s of additional time-out period was given. We gave 4 s extra time for the inter-trial interval before beginning the next trial. Task sessions automatically ended when mice consecutively missed 30 auditory-or visual-Go stimuli. When mice showed more than 80% hit rates for visual and auditory go stimuli, mice were moved on to the third step in the next session.
3. Audiovisual discrimination training: 4 types of stimuli (auditory Go, auditory No-go, visual Go, visual No-go) were pseudo-randomly presented in equal proportion. There was no water reward in the miss trials, and 400 ms of air puff punishment was delivered to the right cheek of mice with the time-out period (6 s) in the FA trials. Task sessions automatically ended when mice consecutively missed 10 auditory-or visual-Go stimuli. If mice achieved correct rates higher than 80% for both auditory and visual discrimination, we considered that the mice learned the discrimination tasks.
4. Treadmill habituation: We first habituated mice to run on the treadmill during the task by presenting the same auditory and visual stimuli when mice ran faster than a speed threshold (3 cm/s). If mice reached an expert level of discrimination (> 80% of correct rates) while showing enough locomotion activity (> 400 trials within 1 hour), mice consecutively performed the task in the stationary block on the immobilized treadmill and in the moving block on the free-rotating treadmill during a single day in random order. If mice showed an expert level of performance in both conditions, they were moved on to the final step. We conducted optic fiber implantation surgery for the optogenetic manipulation experiments when animals passed this step.
5. Audiovisual integration test: auditory, visual, congruent (A_G_V_G_, A_NG_V_NG_), and incongruent (A_G_V_NG_, A_NG_V_G_) stimuli were pseudo-randomly presented in the proportion of 3:3:1:1 or 3:3:2:2 for the main behavioral test (3:3:1:1 for muscimol experiment). 3:3:2:2 (for optogenetic experiments and *in vivo* calcium imaging), or 3:3:1:3 (*in vivo* calcium imaging) ratio was used for the audiovisual integration test to reserve enough trial numbers. There was no water reward and air puff punishment in incongruent trials. Mice conducted both stationary and moving blocks in random order on a single day except for *in vivo* calcium imaging experiments (See the section on *in vivo* calcium imaging experiments).

In moving sessions, we divided trials into two categories: start-moving trials and keep-moving trials. If the average speed before the stimulus onset (-3 ∼ -1.5 s) exceeded the speed threshold (> 3 cm/s), that trial was classified as keep-moving. To evaluate the correlation between locomotion speed and auditory dominance (Supplementary Fig. 1d), we categorized keep-moving incongruent trials according to mean locomotion speed (3 ∼ 9 cm/s, 9 ∼ 15 cm/s, and higher than 15 cm/s) before the stimulus onset (-1 ∼ 0 s). We next determined the auditory dominance rate for each category. To analyze the correlation between trial history and auditory dominance (Supplementary Fig. 1e), we collected and calculated the auditory dominance rates for the following trial sequences: 1) V_G_ or A_NG_ trial followed by A_NG_V_G_ trial, and 2) A_G_ or V_NG_ followed by A_G_V_NG_ trial during the moving sessions. We then estimated the auditory dominance rate based on the modality of the previous trial. To find the relationship between auditory dominance rate and pupil size, we averaged the maximum-normalized pupil size before the stimulus onset (-1 ∼ 0 s) during moving sessions. We then calculated the average value of each variable according to the dominant modality.

#### PPC inactivation by muscimol injection

After the mice became proficient in performing audiovisual discrimination task, we made craniotomies (diameter ∼0.5 mm) on both left and right PPC a day before starting sequential multisensory tests. Except for the injection times, we covered the surface of the craniotomies by applying Dura-Gel (Cambridge NeuroTech) not to damage the cortex. We conducted the sequential multisensory tests as follows: Day 1, test without any injection. Day 2, rest. Day 3, test with muscimol (Sigma-Aldrich) injection. Day 4, rest. Day 5, test with PBS injection. Day 6, rest. Day 7, test with fluorophore-conjugated muscimol (FCM, Invitrogen) injection. We injected muscimol (1 μg/μl; 40 nl in 2 mice or 60nl in 4 mice), PBS (1X; same volume as the muscimol), or 200 nl of FCM into the PPC at both hemispheres (depth 0.35mm) of the head-fixed awake mice at a speed of 1nl/sec. We started the multisensory test 15 minutes after the end of the injection. We ended the behavioral test at the 320^th^ trials (∼45 minutes after the end of the injection) to prevent the effect of overspreading.

#### Optogenetic manipulation

To activate the PV^+^ neurons in the right PPC (Fig. 2d-2j), we delivered a continuous blue light from the laser (1 mW or 5 mW at the fiber tip, 473 nm, Shanghai Laser & Optics Century) starting 200 ms before the stimulus presentation (-0.2 ∼ 1 s from the stimulus onset) in 50% of trials. To inhibit AC_PPC_ neurons in the right AC or AC_STR_ neurons in the bilateral AC (Fig. 6), we delivered a continuous green light from the laser (5 mW, 532 nm, Shanghai Laser & Optics Century) starting 200ms before the stimulus presentation (-0.2 ∼ 1 s from stimulus onset) in 50% of trials. To suppress M2_→AC_ axon terminals in the right AC, we delivered a continuous green light from the laser (5 ∼ 8 mW, 532 nm) starting 200 ms before the stimulus presentation (-0.2 ∼ 1 s from stimulus onset) in 50% of trials. To control the effectiveness of laser light itself during optogenetic inhibition of the target neurons, we conducted control experiments delivering laser lights onto the dental cement.

### Video monitoring of orofacial and body movements of mice performing a task

The data acquisition and analysis were performed as previously reported^86^. We used a camera (G3-GM11-M2020-Genie, Teledyne DALSA) and lens (TEC-M55 2/3" 55mm telecentric lens, C-Mount) to record left eye of task-performing animals at a high resolution (1200 x 800 pixels) by illuminating the left side of the animal’s face with an infrared light source (850 nm, S-IR5850, Skycares). We recorded images at 30 Hz using the Image Acquisition Toolbox provided by MATLAB.

We calculated the pupil size of the left eye of the task-performing mice using the following custom-written MATLAB code: 1) drawing an ROI that covers the left eye, 2) setting the optimal brightness threshold to determine the pupil boundary, 3) converting the ROI image to a binary image by filling in each pixel with the black and white values according to the threshold (*imbinarize*, built-in MATLAB function), 4) filling holes within the pupil boundary to avoid any artifacts from measuring pupil area due to light reflection on the cornea (*imfill*, built-in MATLAB function), 5) identifying a convex hull that includes the inside part of the pupil (*bwconvhull*, a built-in MATLAB function), and 6) counting the number of pixels within the convex hull as the pupil size across frames. We normalized the pupil size using the following equation, resulting in a number ranging from 0 (smallest size) to 1 (largest size):

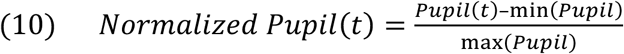

### *In vivo* calcium imaging experiments with miniature microscope

#### Data acquisition

To avoid bleaching of calcium signals during the behavior task, we conducted an imaging experiment in a stationary or moving block one day and tested the other block the next day. Each imaging session was completed when mice showed three consecutive miss trials or when the imaging time reached one hour. We analyzed the same neurons tracked over two consecutive days in imaging data.

We performed an *in vivo* calcium imaging experiment using a miniature fluorescence microscope (Inscopix). Images were acquired at 20 frames per second (10 for AC_PPC_ neurons) using nVista HD software (Inscopix). We cropped superfluous imaging areas that contained no neural signal. The LED power (0.2 ∼ 0.5 mW) and gain value (1 ∼ 4) were adjusted based on the overall brightness of the GCaMP signals. For the simultaneous *in vivo* calcium imaging with optogenetic axon manipulations (Fig. 7 and Fig. 8c and 8d), we used the nVoke system (Inscopix) to acquire images at 10 frames per second with 0.4 mW of red EX-LED light for optogenetic manipulation on ChrimsonR or eNpHR3.0 expressed in axons. The light delivery started 200ms before the stimulus onset and ended at the stimulation offset (-0.2 ∼ 1 s from the stimulus onset).

#### Preprocessing of imaging data

The acquired image data were processed with Inscopix Data Processing Software (IDPS, Inscopix). Images were downsampled by a factor of 4 and spatially filtered (highCutoff: 0.5; lowCutoff: 0.01 for PPC excitatory neurons; highCutoff: 0.5; lowCutoff: 0.005 for the other groups). Filtered images were corrected for lateral motion (IDPS implementation). Motion-corrected images were converted to ΔF/F images where F represents the average intensity of each pixel during the whole imaging time. Principal component analysis -independent component analysis (PCA-ICA) was used to identify cell boundaries on ΔF/F images using IDPS implementation. We next extracted calcium signals by manually inspecting cell boundaries obtained from PCA-ICA on motion-corrected images and exported average intensities within cell boundaries. The same neurons were identified throughout several imaging sessions using longitudinal registration (IDPS implementation), and only neurons that appeared across imaging sessions were selected for further analysis.

#### Calculation of response amplitudes, neuronal type classification, and stimulus selectivity

Exported calcium data were analyzed using custom MATLAB codes. The extracted calcium response data was converted to ΔF/F for each neuron. ΔF/F was calculated as

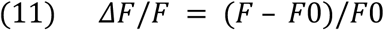

where F0 represents the average fluorescence of a neuron across the imaging time. All the analyses were conducted based on the ΔF/F value.

To determine if a neuron responded to distinct stimuli (Fig. 3c), we compared average calcium activity during baseline (-0.2 ∼ 0 s) and stimulation (0 ∼ 1 s) periods using Wilcoxon signed-rank test (P < 0.01, MATLAB implementation). The response amplitude for each trial was calculated by subtracting the average of baseline activity from the average of calcium activity during the first 1 s after stimulus onset. To dissect neuronal type (Fig. 3c), we compared Go and No-go responses if neurons showed significant responses to Go and No-go stimuli in one modality and classified as Go or No-go neurons if they had a significantly stronger response in the Go or No-go trials. Neurons that showed significant responses in one auditory and one visual stimulus were classified as “Dual,” while neurons with significant responses in three unisensory trial types were classified as “Triple.” “All” type neurons responded significantly in all unisensory trials. To examine how locomotion affects unisensory responses (Supplementary Fig. 2a), we compared the response amplitudes of each neuron from stationary and moving sessions using the Mann-Whitney U-test (P < 0.01).

To calculate the change in spontaneous activity of imaged neurons by locomotion onset and offset, we collected calcium activity and locomotion speed data at all locomotion onsets or offsets (-2 ∼ 2 s) and concatenated them. We then calculated the Pearson correlation coefficient between the two variables. We used the following conditions to define locomotion onset and offset during an imaging session.

Locomotion onset at 0 s must satisfy all of the following conditions:

1. *Average locomotion speed (-1 ∼ -0.5 s) < 0.5cm/s;*
2. *Average locomotion speed (-0.5 ∼ 0 s) < 0.5cm/s;*
3. *Average locomotion speed (0 ∼ 250 ms) > 1cm/s;*
4. *Average locomotion speed (250 ∼ 1000 ms) > 2cm/s;*
5. *No stimulus during -2 ∼ 2 s*.

Locomotion offset at 0 s must satisfy all of the following conditions:

1. *Average locomotion speed (-1 ∼ -0.5 s) > 2 cm/s;*
2. *Average locomotion speed (-0.5 ∼ 0 s) > 1cm/s;*
3. *Average locomotion speed (0 ∼ 250 ms) < 0.5cm/s;*
4. *Average locomotion speed (250 ∼ 1000 ms) < 0.5cm/s;*
5. *No stimulus during -2 ∼ 2 s*.

To calculate the change in M2_AC_ activity at locomotion onset (Fig. 8i), we subtracted mean baseline activity (-1 ∼ -0.5 s from locomotion onset) from mean calcium signal at early (0 ∼ 1.5 s from locomotion onset) or late (3 s from locomotion onset ∼ -1 s from locomotion offset) periods.

To compare the stimulus selectivity (Supplementary Fig. 5), we excluded neurons that were not responsive to the stimulus in either stationary or moving states. We calculated the stimulus selectivity of each neuron as the following formula.

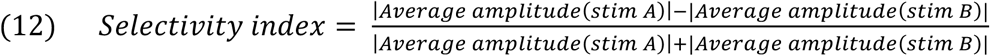

#### Demixed principal component analysis of imaging data

To visualize populational activity during the behavioral task over time, we used demixed principal component analysis (dPCA)^72^. The dPCA was performed using trial-averaged calcium activity aligned at stimulus onset (-1 ∼ 3 s) of all neurons across the eight unisensory trial types (visual hit, visual miss, visual CR, visual FA, auditory hit, auditory miss, auditory CR, and auditory FA). To denoise the calcium data, we temporally downsampled it (factor 2) and applied Gaussian smoothing with a window size of three bins. We excluded sessions that had fewer than three trials of lick or no-lick in unisensory and incongruent trials. The encoding PCs from the dPCA were classified into stimulus-related, decision-related, stimulus-decision interaction-related, and independent axes. To show trajectories for each trial type in Fig. 3g, 4a, 4c, and Supplementary Fig. 6d,7d, we used dPCA results derived from all imaged neurons. For all quantifications, we used the dPCA results from each imaging session. We projected the activities of multisensory trials (congruent and incongruent trials) by using transformation vectors obtained from the dPCA with the unisensory trials.

To quantify the similarities between trial types, we calculated Euclidean distances between coordinates after the stimulus onset (0 ∼ 1 s) in a subspace containing the top 3 stimulus PCs. To compare neural trajectories based on locomotion state, trial-averaged activities of unisensory trials from stationary, start-moving, and keep-moving trials were analyzed by dPCA. To distinguish between start-moving and keep-moving data in the moving sessions, we projected trial data into the encoding PC axes from the dPCA of each moving session. The Euclidean distance between two trajectories was calculated 1 s after the stimulus onset. Projection distance was measured as the Euclidean distance between 0 and 1 s following the stimulus onset.

#### dPCA based classifiers

For classifiers (Fig. 3 and Supplementary Fig. 3,6,7), we projected single-trial activity onto the top encoding PC axes and computed Euclidean distances between single trial data (only correct trials for unisensory classification) and trial-averaged trajectories of unisensory trials (for unisensory classification) or incongruent trials (for incongruent classification) at each time points. Each test trial was classified into a trial type of trial-averaged trajectory that showed the smallest Euclidean distance from the test trial trajectory. If the test trajectory showed a very short projection on the PC axes, its distance to the center of the pre-stimulus trajectories was smaller than its distance to the trial-averaged trajectories at each time point. We counted such trials as incorrect classification.

To determine the significance of similarity with visual trajectories in determining dominant modality (Fig. 4e,f), we first projected the activities of individual unisensory trials onto the top three stimulus encoding PC axes and calculated the Euclidean distance between each trajectory and the trial-averaged target visual trajectory (target visual trajectory: V_G_ for A_NG_V_G_ classification; V_NG_ for A_G_V_NG_ classification). To calculate the distribution of individual visual trials, we used linear discriminant analysis (LDA) to determine an Euclidean distance threshold that distinguishes visual trials from other visual and auditory trials. Finally, we classified individual incongruent trials according to their Euclidean distance to the trial-averaged target visual trajectory, with each trial classified as a visual-dominant trial if its Euclidean distance was less than the threshold. To statistically compare classifier accuracies, we calculated mean classifier accuracy between 0.5 and 1 s following stimulus onset, when the classifiers attained a peak of classification accuracy, and compared between groups.

### *In vivo* calcium imaging experiments with 2-photon microscope

#### Data acquisition

We performed an *in vivo* calcium imaging experiment using a 2-photon microscope (BergamoII, Thorlabs). We adjusted the wavelength of the tuneable excitatory laser (InsightX3, Spectra-Physics) to 920 nm for imaging green channels and 1040 nm for imaging red channels. The laser power was calibrated as 60∼80 mW at the position of the focal plane from the objective lens (XLUMPLFLN20XW, Olympus). We used an ultrasonic clear gel as the immersion medium and covered the gap between the mouse’s head and the objective lens with a fabric curtain to prevent light from the stimulus monitor from being detected. To increase the number of neurons imaged in each session, we conducted three-plane imaging with 30 μm intervals, capturing 7.4 images/s in each plane. The image size was 512 × 512 pixels, corresponding to an actual imaging field size of 600 × 600 μm (imaging center: posterior 3.5 mm and lateral 2.5mm from the bregma). For the red channel imaging to identify VC_PPC_ neurons in each imaging field, we collected images for 1 minute from the same planes we conducted calcium imaging.

#### Processing of imaging data

We used Suite2p for motion correction and cell detection for our 2-photon imaging data^87^. For a brief explanation, we conducted non-rigid motion correction (Suite2p implementation) and used extracted fluorescent signals of soma and neuropil. After cell identification, we manually sorted good cells in each field of view. For the neuropil correction, the corrected fluorescence value for each cell was calculated as follows.

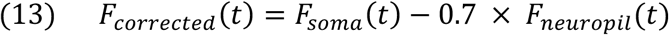

The baseline for ΔF/F calculation was the 10^th^ percentile of *F_corrected_* within a sliding window of 3 mins duration. The rest of the analysis was conducted in the same methods as the data analysis of miniature microscope imaging data. To identify VC_PPC_ neurons in each imaging field, we overlapped the representative red image made by motion correction followed by z-projection to the Suite2p result and manually identified them.

### Histology and Immunohistochemistry

We performed histology experiments to confirm imaging sites after *in vivo* calcium imaging, injection and expression sites of viral vectors, and anatomical circuit tracing data. Mice were anesthetized with avertin (2,2,2-Tribromoethanol (Sigma-Aldrich), 125-250mg/kg, intraperitoneally) and transcardially perfused with 15ml of PBS, followed by 4% paraformaldehyde (PFA, w/v in PBS). Brain samples were post-fixed for 4 hours in the PFA at 4℃, then washed 3 times with PBS for 10 minutes each. After washing, the brain samples were immersed in a 30% sucrose solution (w/v in PBS) for 2-5 days. When the brain samples sank, they were embedded in an optimal cutting temperature medium (Tissue-Tek O.C.T. Compound, Sakura Finetek) and rapidly froze at -80℃. Frozen brain samples were sectioned into 40 μm or 20 μm (for immunostaining) in a coronal direction using a cryocut (Leica). We washed brain slices 3 times in PBS for 10 minutes before mounting them with an anti-bleaching mounting medium containing 4′,6-diamidino-2-phenylindole (DAPI) (Vector Labs). To immunostain the CaMKIIα or PV, we permeabilized the brain sections for 30 min with 0.5% w/v Triton X-100 in PBS, then blocked with 2% w/v normal goat serum (NGS) in PBS for 2 hours at room temperature. The samples were incubated at 4℃ for 48 hours with mouse anti-CaMKIIα antibodies (ab22609, abcam, 1:250 dilution in 2% NGS) or rabbit anti-PV antibodies (PV27, swant, 1:500 dilution in 2% NGS). After treating the primary antibodies, we washed the samples 3 times in PBS for 10 minutes. We used Alexa Fluor 594 goat anti-mouse IgG (A11005, Invitrogen, 1:250 dilution in PBS) to stain CaMKIIα or Alexa Fluor 594 goat anti-rabbit IgG (A21207, Invitrogen, 1:500 dilution in PBS) to stain PV for 2 hours at room temperature. Following that, we washed the samples in PBS 3 times for 10 minutes before mounting them with an anti-bleaching medium with DAPI. Fluorescent images of brain slices were taken using a slide scanner (Zeiss Axio Scan. Z1) or a confocal microscope (Nikon A1 HD25).

### Statistical analysis

We analyzed the data using custom-written codes in MATLAB (Mathworks). All data were presented as mean ± SEM unless otherwise noted. ‘N’ denotes the number of mice or sessions, whereas ‘n’ represents the number of neurons. All statistical analyses were described in the figure legends. We first tested the normality of the data using the Shapiro-Wilk test and applied parametric or non-parametric statistical analyses accordingly. For multiple comparisons, we performed Bonferroni correction on the statistical results or used ANOVA for data that passed the normality test to avoid the increased probability of Type II errors associated with Bonferroni correction when analyzing a high number of pairs (> 6 pairs). All the statistical significances of the datasets were indicated as NS (not significant; *P*≥ 0.1), *#* (*P < 0.1*), * (*P< 0.05*), ** (*P< 0.01*), *** (*P< 0.001*), and **** (*P< 0.0001*).

## Supporting information

Statistical details

## Data availability

Source data are provided with this paper. All the representative data that support the findings are publicly available on GitHub: https://github.com/seungheelee1789/PPC_Choi. Further requests for the data generated in this study can be directed to the corresponding author.

## Code availability

All the custom MATLAB codes to analyze and visualize the data are publicly available on GitHub (https://github.com/seungheelee1789/PPC_Choi).

## Acknowledgements

We thank Dr. Alex C. Kwan for helpful discussions on imaging data analysis and Dr. Jae-Hyun Kim for assistance with pupil data analysis. This work was supported by grants to S.-H.L. from the National Research Foundation of Korea funded by the Ministry of Science & ICT (NRF-2021R1A2C3012159) and the Institute for Basic Science (IBS-R002-A2).

## Author Contributions Statements

I.C. and S.-H.L. conceived and designed all experiments and wrote the manuscript. I.C. performed all experiments and analyzed the data.

## References

1 Stein, B. E. The new handbook of multisensory processing. (MIT Press, 2012).

2 Gielen, S. C., Schmidt, R. A. & Van den Heuvel, P. J. On the nature of intersensory facilitation of reaction time. Percept Psychophys 34, 161–168 (1983). 10.3758/bf03211343

3 Diederich, A. & Colonius, H. Bimodal and trimodal multisensory enhancement: effects of stimulus onset and intensity on reaction time. Percept Psychophys 66, 1388–1404 (2004). 10.3758/bf03195006

4 Raposo, D., Sheppard, J. P., Schrater, P. R. & Churchland, A. K. Multisensory decision-making in rats and humans. J Neurosci 32, 3726–3735 (2012). 10.1523/JNEUROSCI.4998-11.2012

5 Meijer, G. T., Pie, J. L., Dolman, T. L., Pennartz, C. M. A. & Lansink, C. S. Audiovisual Integration Enhances Stimulus Detection Performance in Mice. Front Behav Neurosci 12, 231 (2018). 10.3389/fnbeh.2018.00231

6 Colavita, F. B. Human Sensory Dominance. Perception & Psychophysics 16, 409–412 (1974). 10.3758/Bf03203962

7 McGurk, H. & MacDonald, J. Hearing lips and seeing voices. Nature 264, 746–748 (1976). 10.1038/264746a0

8 Battaglia, P. W., Jacobs, R. A. & Aslin, R. N. Bayesian integration of visual and auditory signals for spatial localization. J Opt Soc Am A Opt Image Sci Vis 20, 1391–1397 (2003). 10.1364/josaa.20.001391

9 Alais, D. & Burr, D. The ventriloquist effect results from near-optimal bimodal integration. Curr Biol 14, 257–262 (2004). 10.1016/j.cub.2004.01.029

10 Fetsch, C. R., Turner, A. H., DeAngelis, G. C. & Angelaki, D. E. Dynamic reweighting of visual and vestibular cues during self-motion perception. J Neurosci 29, 15601–15612 (2009). 10.1523/JNEUROSCI.2574-09.2009

11 Fetsch, C. R., Pouget, A., DeAngelis, G. C. & Angelaki, D. E. Neural correlates of reliability-based cue weighting during multisensory integration. Nat Neurosci 15, 146–154 (2011). 10.1038/nn.2983

12 Song, Y. H. et al. A Neural Circuit for Auditory Dominance over Visual Perception. Neuron 93, 940-+ (2017). 10.1016/j.neuron.2017.01.006

13 Choi, I. & Lee, S. H. Reduced activity of parvalbumin-positive interneurons in the posterior parietal cortex causes visually dominant multisensory decisions in freely navigating mice. Mol Brain 15, 82 (2022). 10.1186/s13041-022-00968-x

14 Ernst, M. O. & Banks, M. S. Humans integrate visual and haptic information in a statistically optimal fashion. Nature 415, 429–433 (2002). 10.1038/415429a

15 Kording, K. P. & Wolpert, D. M. Bayesian integration in sensorimotor learning. Nature 427, 244–247 (2004). 10.1038/nature02169

16 Coen, P., Sit, T. P. H., Wells, M. J., Carandini, M. & Harris, K. D. Mouse frontal cortex mediates additive multisensory decisions. Neuron 111, 2432–2447 e2413 (2023). 10.1016/j.neuron.2023.05.008

17 Keil, J., Muller, N., Ihssen, N. & Weisz, N. On the variability of the McGurk effect: audiovisual integration depends on prestimulus brain states. Cereb Cortex 22, 221–231 (2012). 10.1093/cercor/bhr125

18 Jiang, W., Jiang, H. & Stein, B. E. Two corticotectal areas facilitate multisensory orientation behavior. J Cogn Neurosci 14, 1240–1255 (2002). 10.1162/089892902760807230

19 Avillac, M., Ben Hamed, S. & Duhamel, J. R. Multisensory integration in the ventral intraparietal area of the macaque monkey. J Neurosci 27, 1922–1932 (2007). 10.1523/JNEUROSCI.2646-06.2007

20 Mallick, D. B., Magnotti, J. F. & Beauchamp, M. S. Variability and stability in the McGurk effect: contributions of participants, stimuli, time, and response type. Psychon Bull Rev 22, 1299–1307 (2015). 10.3758/s13423-015-0817-4

21 Sinnett, S., Spence, C. & Soto-Faraco, S. Visual dominance and attention: the Colavita effect revisited. Percept Psychophys 69, 673–686 (2007). 10.3758/bf03193770

22 Crochet, S., Lee, S. H. & Petersen, C. C. H. Neural Circuits for Goal-Directed Sensorimotor Transformations. Trends Neurosci 42, 66–77 (2019). 10.1016/j.tins.2018.08.011

23 Esmaeili, V. et al. Rapid suppression and sustained activation of distinct cortical regions for a delayed sensory-triggered motor response. Neuron 109, 2183–2201 e2189 (2021). 10.1016/j.neuron.2021.05.005

24 Choi, I., Lee, J. Y. & Lee, S. H. Bottom-up and top-down modulation of multisensory integration. Curr Opin Neurobiol 52, 115–122 (2018). 10.1016/j.conb.2018.05.002

25 McGinley, M. J. et al. Waking State: Rapid Variations Modulate Neural and Behavioral Responses. Neuron 87, 1143–1161 (2015). 10.1016/j.neuron.2015.09.012

26 Ayaz, A., Saleem, A. B., Scholvinck, M. L. & Carandini, M. Locomotion controls spatial integration in mouse visual cortex. Curr Biol 23, 890–894 (2013). 10.1016/j.cub.2013.04.012

27 Bennett, C., Arroyo, S. & Hestrin, S. Subthreshold Mechanisms Underlying State-Dependent Modulation of Visual Responses. Neuron 80, 350–357 (2013). 10.1016/j.neuron.2013.08.007

28 Dipoppa, M. et al. Vision and Locomotion Shape the Interactions between Neuron Types in Mouse Visual Cortex. Neuron 98, 602-+ (2018). 10.1016/j.neuron.2018.03.037

29 Fu, Y. et al. A Cortical Circuit for Gain Control by Behavioral State. Cell 156, 1139–1152 (2014). 10.1016/j.cell.2014.01.050

30 Lee, A. M. et al. Identification of a Brainstem Circuit Regulating Visual Cortical State in Parallel with Locomotion. Neuron 83, 455–466 (2014). 10.1016/j.neuron.2014.06.031

31 Niell, C. M. & Stryker, M. P. Modulation of Visual Responses by Behavioral State in Mouse Visual Cortex. Neuron 65, 472–479 (2010). 10.1016/j.neuron.2010.01.033

32 Paken, J. M. P. et al. Behavioral-state modulation of inhibition is context-dependent and cell type specific in mouse visual cortex. Elife 5 (2016). ARTN e14985 10.7554/eLife.14985

33 Polack, P. O., Friedman, J. & Golshani, P. Cellular mechanisms of brain state-dependent gain modulation in visual cortex. Nature Neuroscience 16, 1331–U1227 (2013). 10.1038/nn.3464

34 Vinck, M., Batista-Brito, R., Knoblich, U. & Cardin, J. A. Arousal and Locomotion Make Distinct Contributions to Cortical Activity Patterns and Visual Encoding. Neuron 86, 740–754 (2015). 10.1016/j.neuron.2015.03.028

35 McGinley, M. J., David, S. V. & McCormick, D. A. Cortical Membrane Potential Signature of Optimal States for Sensory Signal Detection. Neuron 87, 179–192 (2015). 10.1016/j.neuron.2015.05.038

36 Zhou, M. et al. Scaling down of balanced excitation and inhibition by active behavioral states in auditory cortex. Nature Neuroscience 17, 841–850 (2014). 10.1038/nn.3701

37 Nelson, A. et al. A Circuit for Motor Cortical Modulation of Auditory Cortical Activity. Journal of Neuroscience 33, 14342-+ (2013). 10.1523/Jneurosci.2275-13.2013

38 Schneider, D. M., Nelson, A. & Mooney, R. A synaptic and circuit basis for corollary discharge in the auditory cortex. Nature 513, 189-+ (2014). 10.1038/nature13724

39 Schneider, D. M., Sundararajan, J. & Mooney, R. A cortical filter that learns to suppress the acoustic consequences of movement. Nature 561, 391-+ (2018). 10.1038/s41586-018-0520-5

40 Runyan, C. A., Piasini, E., Panzeri, S. & Harvey, C. D. Distinct timescales of population coding across cortex. Nature 548, 92–96 (2017). 10.1038/nature23020

41 Williamson, R. S., Hancock, K. E., Shinn-Cunningham, B. G. & Polley, D. B. Locomotion and Task Demands Differentially Modulate Thalamic Audiovisual Processing during Active Search. Curr Biol 25, 1885–1891 (2015). 10.1016/j.cub.2015.05.045

42 Zhou, Y. & Freedman, D. J. Posterior parietal cortex plays a causal role in perceptual and categorical decisions. Science 365, 180–185 (2019). 10.1126/science.aaw8347

43 Lyamzin, D. & Benucci, A. The mouse posterior parietal cortex: Anatomy and functions. Neurosci Res 140, 14–22 (2019). 10.1016/j.neures.2018.10.008

44 Freedman, D. J. & Ibos, G. An Integrative Framework for Sensory, Motor, and Cognitive Functions of the Posterior Parietal Cortex. Neuron 97, 1219–1234 (2018). 10.1016/j.neuron.2018.01.044

45 Goard, M. J., Pho, G. N., Woodson, J. & Sur, M. Distinct roles of visual, parietal, and frontal motor cortices in memory-guided sensorimotor decisions. Elife 5 (2016). 10.7554/eLife.13764

46 Nikbakht, N., Tafreshiha, A., Zoccolan, D. & Diamond, M. E. Supralinear and Supramodal Integration of Visual and Tactile Signals in Rats: Psychophysics and Neuronal Mechanisms. Neuron 97, 626–639 e628 (2018). 10.1016/j.neuron.2018.01.003

47 Oude Lohuis, M. N., Marchesi, P., Pennartz, C. M. A. & Olcese, U. Functional (ir)relevance of posterior parietal cortex during audiovisual change detection. J Neurosci 42, 5229–5245 (2022). 10.1523/JNEUROSCI.2150-21.2022

48 Licata, A. M. et al. Posterior Parietal Cortex Guides Visual Decisions in Rats. J Neurosci 37, 4954–4966 (2017). 10.1523/JNEUROSCI.0105-17.2017

49 Morcos, A. S. & Harvey, C. D. History-dependent variability in population dynamics during evidence accumulation in cortex. Nat Neurosci 19, 1672–1681 (2016). 10.1038/nn.4403

50 Scott, B. B. et al. Fronto-parietal Cortical Circuits Encode Accumulated Evidence with a Diversity of Timescales. Neuron 95, 385–398 e385 (2017). 10.1016/j.neuron.2017.06.013

51 Hwang, E. J., Dahlen, J. E., Mukundan, M. & Komiyama, T. History-based action selection bias in posterior parietal cortex. Nat Commun 8, 1242 (2017). 10.1038/s41467-017-01356-z

52 Driscoll, L. N., Pettit, N. L., Minderer, M., Chettih, S. N. & Harvey, C. D. Dynamic Reorganization of Neuronal Activity Patterns in Parietal Cortex. Cell 170, 986–999 e916 (2017). 10.1016/j.cell.2017.07.021

53 Harvey, C. D., Coen, P. & Tank, D. W. Choice-specific sequences in parietal cortex during a virtual-navigation decision task. Nature 484, 62–68 (2012). 10.1038/nature10918

54 Pho, G. N., Goard, M. J., Woodson, J., Crawford, B. & Sur, M. Task-dependent representations of stimulus and choice in mouse parietal cortex. Nat Commun 9, 2596 (2018). 10.1038/s41467-018-05012-y

55 Tseng, S. Y., Chettih, S. N., Arlt, C., Barroso-Luque, R. & Harvey, C. D. Shared and specialized coding across posterior cortical areas for dynamic navigation decisions. Neuron 110, 2484–2502 e2416 (2022). 10.1016/j.neuron.2022.05.012

56 Raposo, D., Kaufman, M. T. & Churchland, A. K. A category-free neural population supports evolving demands during decision-making. Nature Neuroscience 17, 1784–1792 (2014). 10.1038/nn.3865

57 Akrami, A., Kopec, C. D., Diamond, M. E. & Brody, C. D. Posterior parietal cortex represents sensory history and mediates its effects on behaviour. Nature 554, 368–372 (2018). 10.1038/nature25510

58 Zhong, L. et al. Causal contributions of parietal cortex to perceptual decision-making during stimulus categorization. Nat Neurosci 22, 963–973 (2019). 10.1038/s41593-019-0383-6

59 Hanks, T. D. et al. Distinct relationships of parietal and prefrontal cortices to evidence accumulation. Nature 520, 220–223 (2015). 10.1038/nature14066

60 Erlich, J. C., Brunton, B. W., Duan, C. A., Hanks, T. D. & Brody, C. D. Distinct effects of prefrontal and parietal cortex inactivations on an accumulation of evidence task in the rat. Elife 4 (2015). 10.7554/eLife.05457

61 Gallero-Salas, Y. et al. Sensory and Behavioral Components of Neocortical Signal Flow in Discrimination Tasks with Short-Term Memory. Neuron 109, 135–148 e136 (2021). 10.1016/j.neuron.2020.10.017

62 Yao, J. D., Gimoto, J., Constantinople, C. M. & Sanes, D. H. Parietal Cortex Is Required for the Integration of Acoustic Evidence. Curr Biol 30, 3293–3303 e3294 (2020). 10.1016/j.cub.2020.06.017

63 Zingg, B. et al. Neural networks of the mouse neocortex. Cell 156, 1096–1111 (2014). 10.1016/j.cell.2014.02.023

64 Zhang, S. et al. Organization of long-range inputs and outputs of frontal cortex for top-down control. Nat Neurosci 19, 1733–1742 (2016). 10.1038/nn.4417

65 Lewis, J. W. & Van Essen, D. C. Corticocortical connections of visual, sensorimotor, and multimodal processing areas in the parietal lobe of the macaque monkey. J Comp Neurol 428, 112–137 (2000). 10.1002/1096-9861(20001204)428:1<112::aid-cne8>3.0.co;2-9

66 Driver, J. & Noesselt, T. Multisensory interplay reveals crossmodal influences on ’sensory-specific’ brain regions, neural responses, and judgments. Neuron 57, 11–23 (2008). 10.1016/j.neuron.2007.12.013

67 Schlack, A., Sterbing-D’Angelo, S. J., Hartung, K., Hoffmann, K. P. & Bremmer, F. Multisensory space representations in the macaque ventral intraparietal area. J Neurosci 25, 4616–4625 (2005). 10.1523/JNEUROSCI.0455-05.2005

68 Olcese, U., Iurilli, G. & Medini, P. Cellular and synaptic architecture of multisensory integration in the mouse neocortex. Neuron 79, 579–593 (2013). 10.1016/j.neuron.2013.06.010

69 Wallace, M. T., Ramachandran, R. & Stein, B. E. A revised view of sensory cortical parcellation. Proc Natl Acad Sci U S A 101, 2167–2172 (2004). 10.1073/pnas.0305697101

70 Shimaoka, D., Harris, K. D. & Carandini, M. Effects of Arousal on Mouse Sensory Cortex Depend on Modality (vol 22, pg 3160, 2018). Cell Reports 25, 3230–3230 (2018). 10.1016/j.celrep.2018.11.105

71 Schneider, D. M. & Mooney, R. Motor-related signals in the auditory system for listening and learning. Curr Opin Neurobiol 33, 78–84 (2015). 10.1016/j.conb.2015.03.004

72 Kobak, D. et al. Demixed principal component analysis of neural population data. Elife 5 (2016). 10.7554/eLife.10989

73 Chen, L., Wang, X. X., Ge, S. Y. & Xiong, Q. J. Medial geniculate body and primary auditory cortex differentially contribute to striatal sound representations. Nature Communications 10 (2019). ARTN 418 10.1038/s41467-019-08350-7

74 Guo, L., Walker, W. I., Ponvert, N. D., Penix, P. L. & Jaramillo, S. Stable representation of sounds in the posterior striatum during flexible auditory decisions. Nature Communications 9 (2018). ARTN 1534 10.1038/s41467-018-03994-3

75 Xiong, Q. J., Znamenskiy, P. & Zador, A. M. Selective corticostriatal plasticity during acquisition of an auditory discrimination task. Nature 521, 348-+ (2015). 10.1038/nature14225

76 Ponvert, N. D. & Jaramillo, S. Auditory Thalamostriatal and Corticostriatal Pathways Convey Complementary Information about Sound Features. Journal of Neuroscience 39, 271–280 (2019). 10.1523/Jneurosci.1188-18.2018

77 Lin, P. A., Asinof, S. K., Edwards, N. J. & Isaacson, J. S. Arousal regulates frequency tuning in primary auditory cortex. Proc Natl Acad Sci U S A 116, 25304–25310 (2019). 10.1073/pnas.1911383116

78 Welch, R. B. & Warren, D. H. Immediate perceptual response to intersensory discrepancy. Psychol Bull 88, 638–667 (1980).

79 Repp, B. H. & Penel, A. Auditory dominance in temporal processing: New evidence from synchronization with simultaneous visual and auditory sequences. J Exp Psychol Human 28, 1085–1099 (2002). 10.1037//0096-1523.28.5.1085

80 Crapse, T. B. & Sommer, M. A. Corollary discharge across the animal kingdom. Nat Rev Neurosci 9, 587–600 (2008). 10.1038/nrn2457

81 Kuan, A. T. et al. Synaptic wiring motifs in posterior parietal cortex support decision-making. bioRxiv, 2022.2004.2013.488176 (2022). 10.1101/2022.04.13.488176

82 Romanski, L. M. Representation and integration of auditory and visual stimuli in the primate ventral lateral prefrontal cortex. Cereb Cortex 17 **Suppl 1**, i61–69 (2007). 10.1093/cercor/bhm099

83 Calvert, G. A., Campbell, R. & Brammer, M. J. Evidence from functional magnetic resonance imaging of crossmodal binding in the human heteromodal cortex. Curr Biol 10, 649–657 (2000). 10.1016/s0960-9822(00)00513-3

84 Gulati, S., Cao, V. Y. & Otte, S. Multi-layer Cortical Ca2+ Imaging in Freely Moving Mice with Prism Probes and Miniaturized Fluorescence Microscopy. Jove-J Vis Exp (2017). ARTN e55579 10.3791/55579

85 Goldey, G. J. et al. Removable cranial windows for long-term imaging in awake mice. Nat Protoc 9, 2515–2538 (2014). 10.1038/nprot.2014.165

86 Kim, J. H., Ma, D. H., Jung, E., Choi, I. & Lee, S. H. Gated feedforward inhibition in the frontal cortex releases goal-directed action. Nat Neurosci 24, 1452–1464 (2021). 10.1038/s41593-021-00910-9

87 Pachitariu, M. et al. Suite2p: beyond 10,000 neurons with standard two-photon microscopy. bioRxiv, 061507 (2017). 10.1101/061507

