## Supplementary figures and captions for "Locomotion-dependent auditory gating to the parietal cortex guides multisensory decisions"

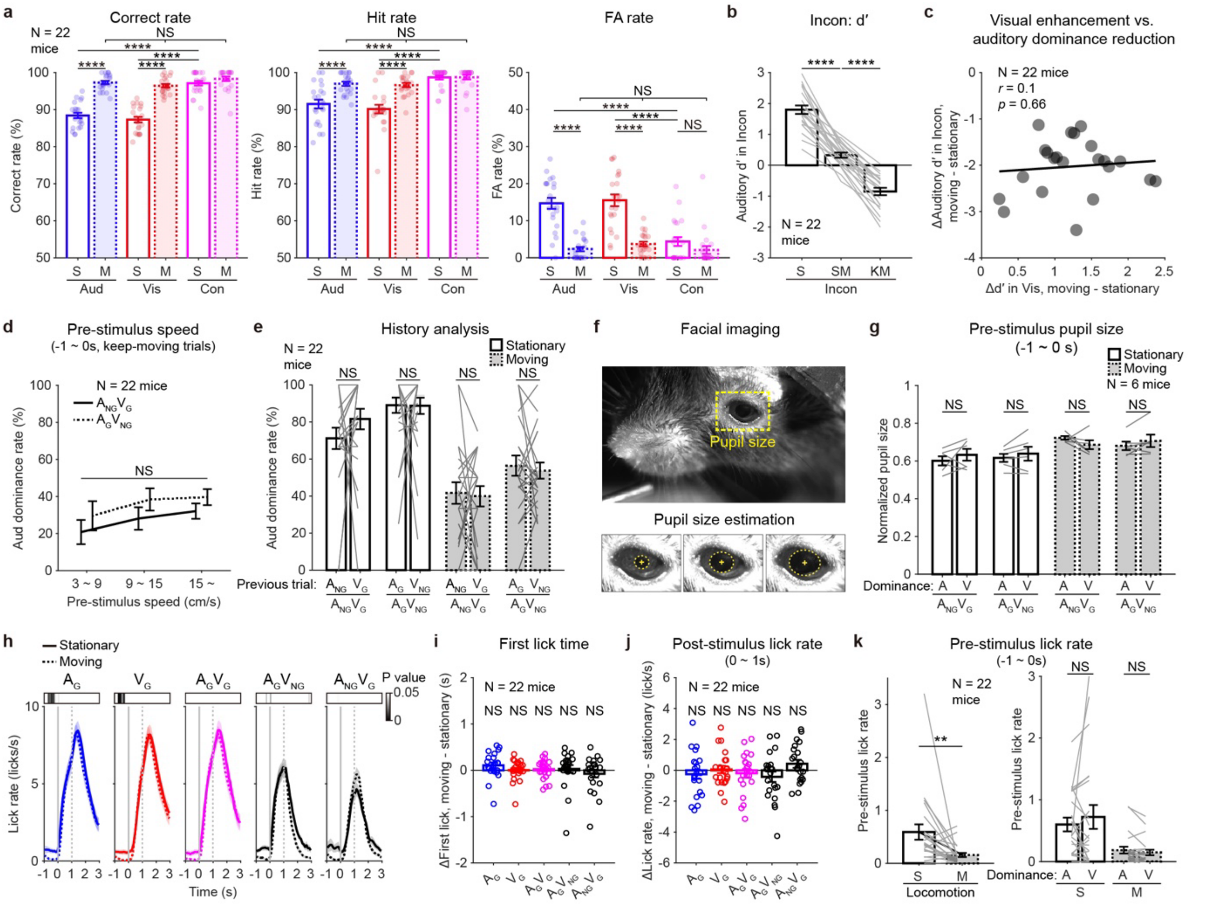


**Supplementary Fig. 1 | Behavioral performance analysis of mice under the audiovisual discrimination task.**

**a**, Behavioral performances in correct rates (left), hit rates (middle), and FA rates (right) across the trial types during the stationary (S) and moving (M) sessions.

**b**, Auditory d′ in incongruent trials across the locomotion states. S, stationary; SM, start-moving; KM, keep-moving. Gray lines represent individual mouse data.

**c**, Correlation between the enhancement in visual discrimination and the reduction of auditory dominance during locomotion. *r*, Pearson correlation coefficient; *p*, significance of the Pearson correlation. Black line, linear regression (slope = 0.11).

**d**, Auditory dominance rates at different pre-stimulus locomotion speeds in $A_{NG}V_{G}$ (gray solid line), $A_{G}V_{NG}$ (gray dashed line), and total incongruent trials (black) from the keep-moving state.

**e**, Auditory dominance rates in incongruent trials after different types of unisensory trials.

**f**, Example snapshot of a mouse head on the treadmill (top) and automatically estimated pupil boundaries at three different time points (bottom).

**g**, Mean pre-stimulus pupil sizes in incongruent trials when mice showed auditory (A) or visual (V) dominance at different locomotion states.

**h**, Lick rates aligned to the stimulus onset in the different trial types with go stimuli under different locomotion states. Solid lines represent stationary session; Dotted lines represent moving session. Vertical gray lines, stimulus onset (solid) and offset (dotted). P values from Wilcoxon signed-rank test with Bonferroni correction for each time point were indicated.

**i**, Changes in trial-average first lick time after the stimulus onset during moving sessions compared to stationary sessions.

**j**, Changes in average post-stimulus lick rates during moving sessions compared to stationary sessions.

**k**, Left, mean pre-stimulus lick rates from all trials during stationary and moving sessions. Right, average pre-stimulus lick rates in incongruent trials according to dominant modalities and locomotion states.

Data are presented as means ± SEM. Circles and gray lines are individual mouse data. Source data are available in Source Data File. Sample numbers and statistical information are listed in Supplementary Data 1. NS (not significant), $**P < 0.01$, $****P < 0.0001$.

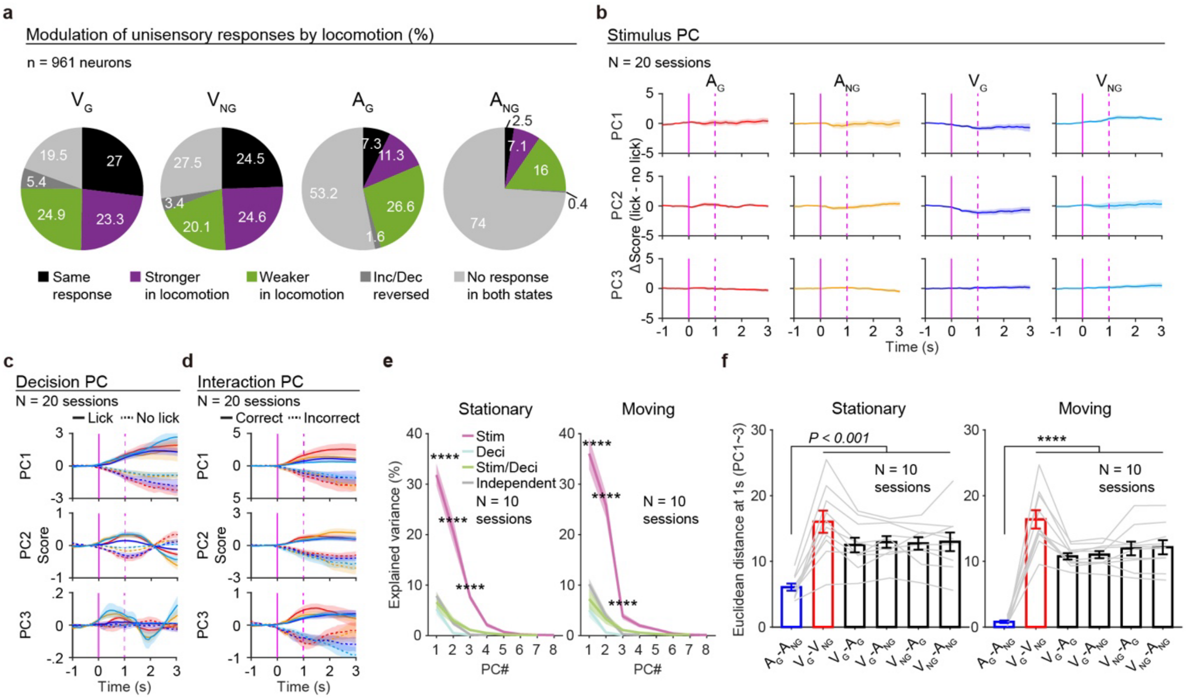


**Supplementary Fig. 2 | Unisensory response changes during locomotion and additional dPCA analysis of the PPC excitatory neurons.**

**a**, Changes in unisensory responses of the PPC excitatory neurons by locomotion.

**b**, Differences in the top 3 PC values in lick trials compared to no-lick trials in the stimulus PC axes after dPCA (N = 20 sessions; 10 stationary sessions; 10 moving sessions). Note that the PCs in the stimulus axes were not different between lick and no-lick decisions of mice (Δ Score ≈ 0). Color, unisensory trial type. Magenta, stimulus onset (solid) and offset (dotted).

**c**, Comparisons of the top 3 PC values in the decision axes. Colors indicate unisensory types as shown in **b**. Note that decision PCs clearly dissect lick (solid line) and no-lick (dotted line) trials.

**d**, Same as **c**, but for the PCs in the interaction axes. Note that interaction PCs clearly dissect correct and incorrect trials

**e**, Percentage of explained variances of top 20 demixed PCs from dPCA of the stationary (left) and the moving sessions (right). Asterisks indicate statistical comparisons between the stimulus PCs and other PCs.

**f**, Mean Euclidean distances in a pair of the unisensory trajectories in the stimulus subspace isolated from stationary (left) and moving (right) sessions. Note that the Euclidean distances are larger between visual Go and No-go trajectories than those between auditory Go and No-go trajectories in both conditions. Gray lines indicate individual data.

Data are presented as means ± SEM. Source data are available in Source Data File. Sample numbers and statistical information are listed in Supplementary Data 1. NS, not significant; *****P* < 0.001.

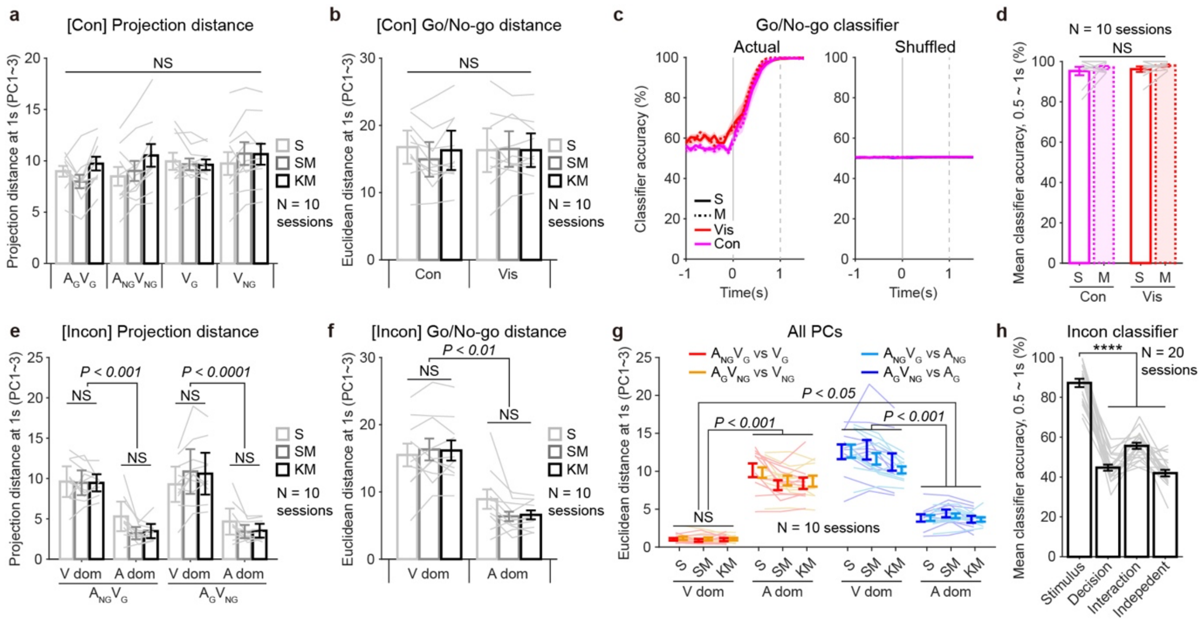


**Supplementary Fig. 3 | Additional dPCA results from the PPC neurons in the multisensory trials.**

**a**, Projection distances of neural trajectories in the stimulus subspace during congruent audiovisual and visual trials at different states. S, stationary; SM, start-moving; KM, keep-moving.

**b**, Euclidean distances between stimulus trajectories of Go and No-go trials (Go/No-go distance) in congruent (Con) and visual (Vis) trials at different states.

**c**, Classification accuracies of the visual Go and No-go trials and congruent Go and No-go trials in the stationary (S) and the moving (M) states using actual data (left) and shuffled data (right) in trial labels. Solid lines mean accuracies in stationary sessions; Dashed lines mean accuracies in moving sessions. For shuffled data, lines represent upper boundaries of 95% confidence intervals in the accuracies from 500 repetitions. Vertical gray lines, stimulus onset (solid) and offset (dashed).

**d**, Mean Go/No-go classifier accuracies during the 0.5 ~ 1 s period after the stimulus onset.

**e**,**f**, Same as **a**,**b**, but for incongruent audiovisual trials. Note that the incongruent trials were further sorted into visual-dominant and auditory-dominant trials.

**g**, Euclidean distances between the incongruent audiovisual trajectories and the unisensory visual or auditory trajectories of all PCs in the stimulus subspace.

**h**, Mean classifier accuracies in identifying the four types of incongruent trials (two stimulus conditions with visual- or auditory-dominant decisions) by using neural trajectories in the stimulus, decision, interaction, and independent subspaces.

Data are presented as means ± SEM. Connected lines indicate individual data. Source data are available in Source Data File. Sample numbers and statistical information are listed in Supplementary Data 1. NS, not significant; ****$P < 0.0001$.

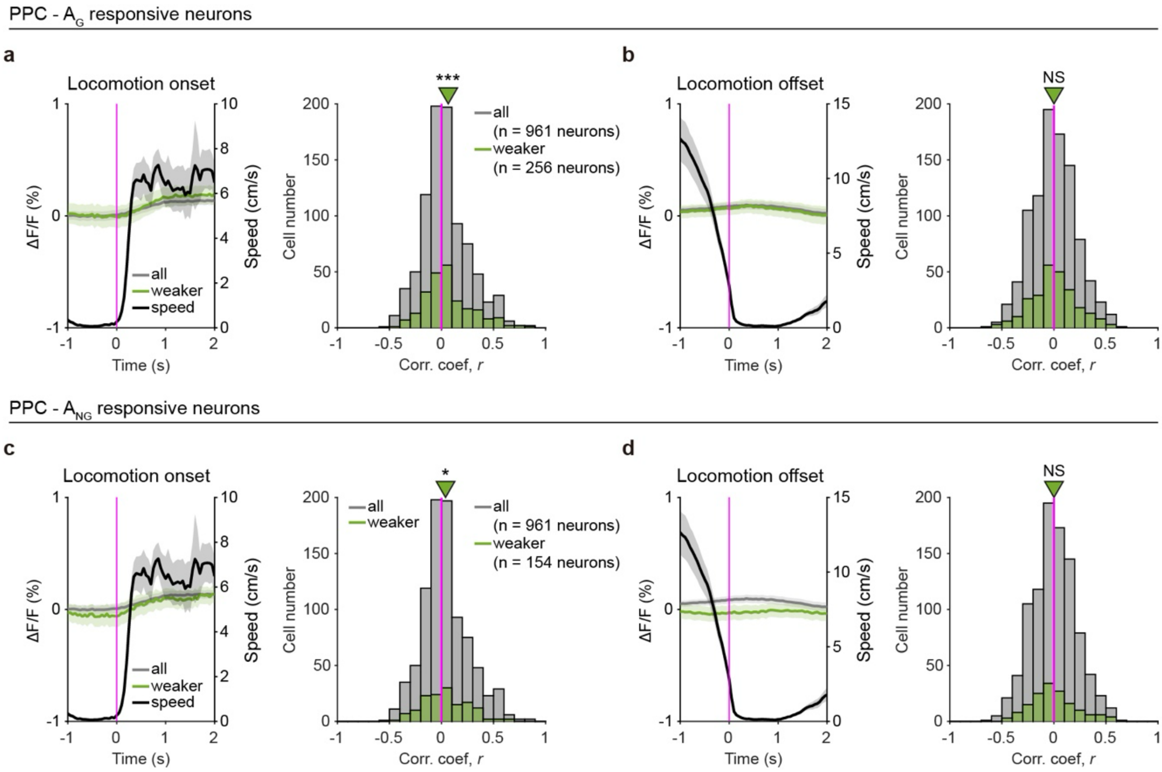


**Supplementary Fig. 4 | Locomotion-related activities in the PPC excitatory neurons.**

**a**, Left, the mean calcium trace of all PPC neurons (gray) and $A_G$ neurons (green) that showed weaker auditory responses during locomotion (as shown in Supplementary Data Fig. 2a), along with the mean locomotion speed of mice (black) aligned to the locomotion onset (magenta). Right, a histogram of Pearson correlation coefficients between the activity changes in those neurons and the locomotion speeds during -2 ~ 2 s of the locomotion onset. Magenta, zero line. Arrowheads, mean correlation coefficients.

**b**, Same as **a**, but for the locomotion offset.

**c**,**d**, Same as **a**,**b**, but for the $A_{NG}$ neurons (green) that showed weaker auditory responses during locomotion.

Traces with shades are means ± SEM. Source data are available in Source Data File. Sample numbers and statistical information are listed in Supplementary Data 1. NS, not significant; *$P$ < 0.05; ***$P$ < 0.001.

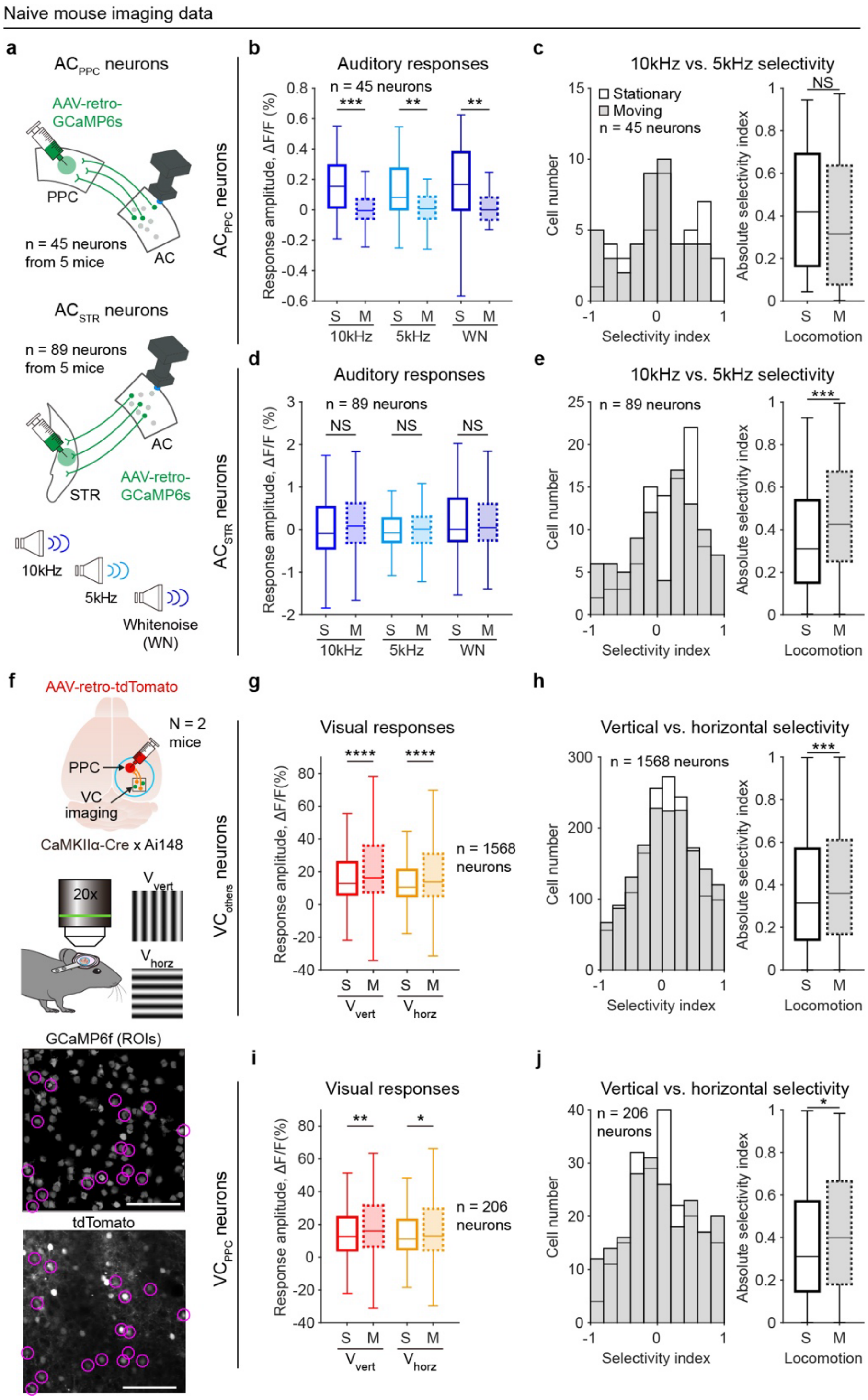


**Supplementary Fig. 5 | Modulation of sensory encoding in the AC and VC by locomotion.**

**a**, Schematic diagram of *in vivo* calcium imaging of $AC_{PPC}$ neurons (top), $AC_{STR}$ neurons (middle), and the auditory stimuli used for the imaging (bottom).

**b**, Auditory responses of $AC_{PPC}$ neurons according to locomotion states. S, stationary condition; M, moving condition.

**c**, Left, stimulus selectivity for two pure tones in stationary and moving sessions from the same $AC_{PPC}$ neurons. Right, mean absolute selectivity index of all $AC_{PPC}$ neurons in stationary and moving sessions.

**d** and **e**, Same for **b** and **c**, but for $AC_{STR}$ neurons.

**f**, Top, schematic illustration of *in vivo* 2-photon calcium imaging in the VC while identifying PPC-projecting neurons. Bottom, identified $VC_{PPC}$ neurons (magenta circles) in an example field of view. Scale bar, 100μm.

**g**, Visual responses of VC neurons that did not project to the PPC, according to locomotion states.

**h,** Left, stimulus selectivity for two drifting gratings in stationary and moving sessions from the same neurons. Right, mean absolute selectivity index in stationary and moving sessions.

**i** and **j**, Same for **g** and **h**, but for the $VC_{PPC}$ neurons.

Data are presented as means ± SEM. For box plots, central lines indicate medians; boxes represent the $25^{th}$-$75^{th}$ percentiles; whiskers show maximum/minimum values excluding outliers. Source data are available in Source Data File. Sample numbers and statistical information are listed in Supplementary Data 1. NS, not significant; **P* < 0.05, ***P* < 0.05, ****P* < 0.001, *****P* < 0.0001.

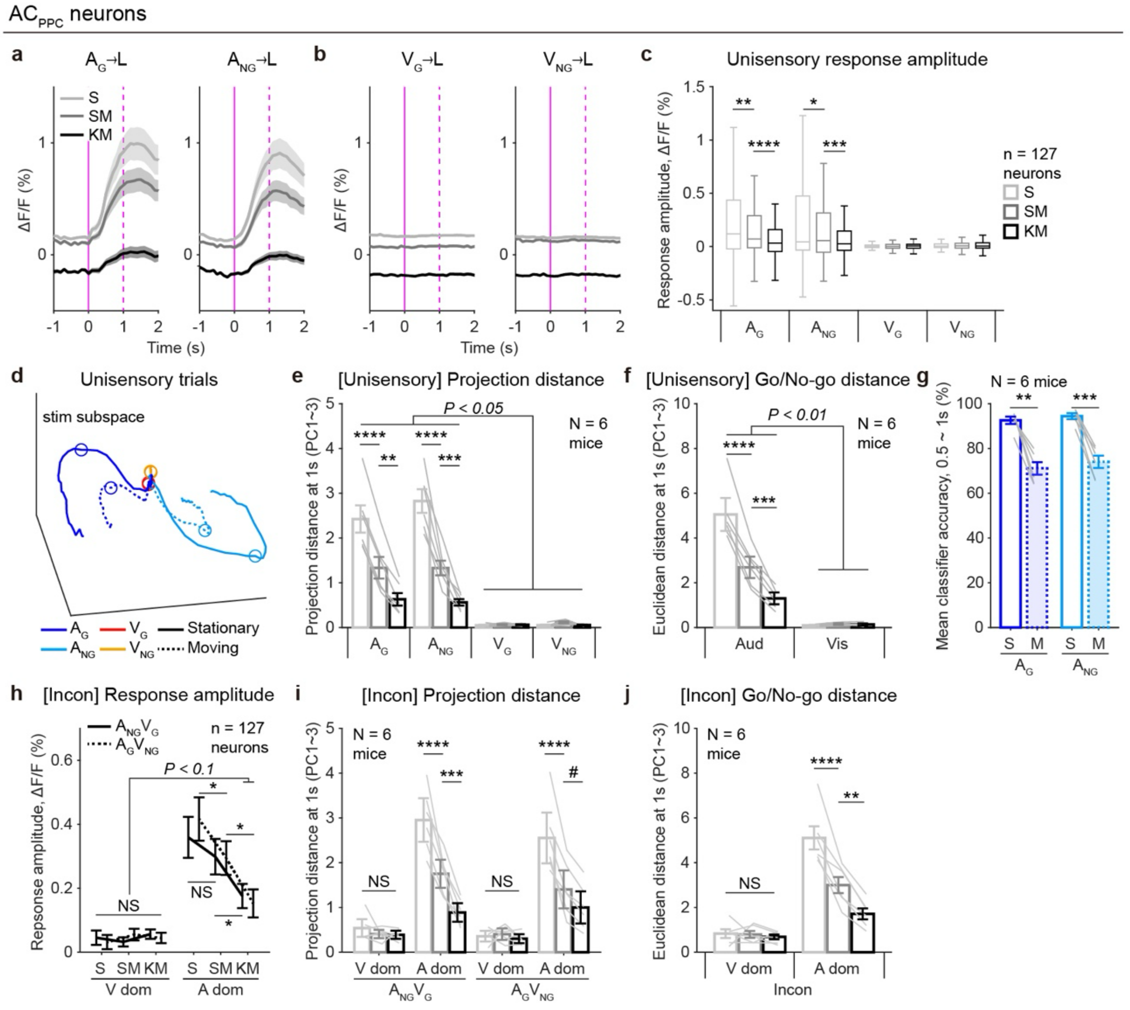


**Supplementary Fig. 6 | Neural activities of the $AC_{PPC}$ neurons at different locomotion states.**

**a**, Mean calcium traces of the $AC_{PPC}$ neurons in the trials with the $A_G$ stimulus (left) and with the $A_{NG}$ stimulus (right) at different locomotion states. Note that the responses were suppressed when mice showed longer locomotion. Magenta, stimulus onset (solid) and offset (dotted). S, stationary; SM, start-moving; KM, keep-moving.

**b**, Same as **a**, but for the $AC_{PPC}$ neurons in the trials with the $V_G$ (left) and $V_{NG}$ stimuli (right) at different locomotion states.

**c**, Response amplitudes of the $AC_{PPC}$ neurons in the unisensory trials at different locomotion states.

**d**, Neural trajectories of the $AC_{PPC}$ neurons in the top three PCs of the stimulus subspace from the dPCA. Four colors in trajectories represent unisensory trial types. Solid lines, stationary conditions; dotted lines, moving conditions.

**e**,**f**, Projections distances of the unisensory trajectories (**e**) and distances between Go and No-go trajectories (**f**) at different locomotion states.

**g**, Mean classifier accuracies of auditory trials during 0.5 ~ 1 s post-stimulus period. S, stationary condition; M, moving condition.
**h**-**j**, Same as **c**,**e**,**f**, but for incongruent trials.
Data are presented as means ± SEM. Overlaid gray lines indicate individual data. For box plots, central lines indicate medians; boxes represent the $25^{th}$-$75^{th}$ percentiles; whiskers show maximum/minimum values excluding outliers. Source data are available in Source Data File. Sample numbers and statistical information are listed in Supplementary Data 1. NS, not significant; #$P < 0.1$, *$P < 0.05$, **$P < 0.01$, ***$P < 0.001$, ****$P < 0.0001$.

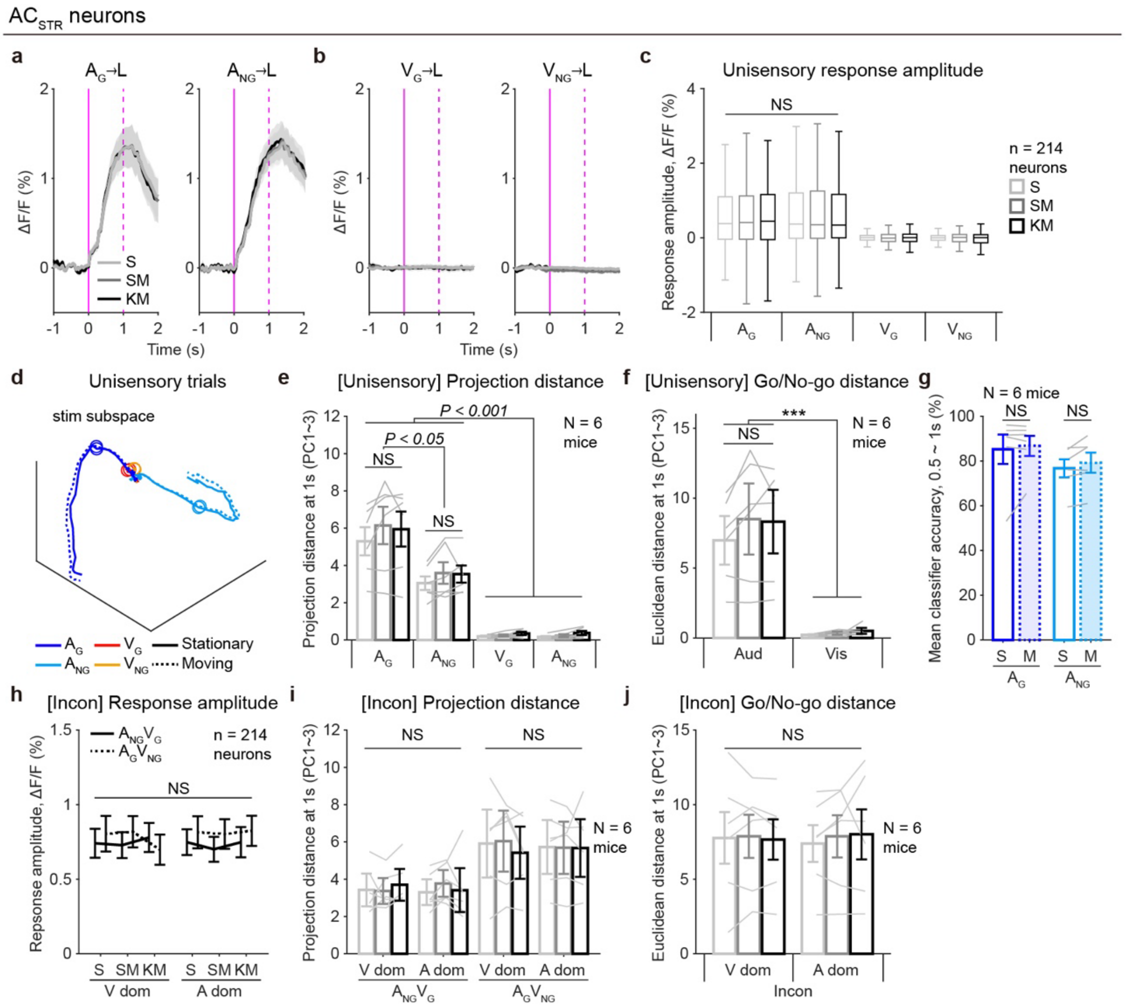


**Supplementary Fig. 7 | Neural activities of the $AC_{STR}$ neurons at different locomotion states.**

**a**-**j**, Same as Supplementary Fig. 6, but for the $AC_{STR}$ neurons.

**a**, Mean calcium traces of the $AC_{STR}$ neurons in the trials with the $A_G$ stimulus (left) and with the $A_{NG}$ stimulus (right) at different locomotion states. Magenta, stimulus onset (solid) and offset (dotted). S, stationary; SM, start-moving; KM, keep-moving.

**b**, Same as **a**, but for the $AC_{STR}$ neurons in the trials with the $V_G$ (left) and $V_{NG}$ stimuli (right) at different locomotion states.

**c**, Response amplitudes of the $AC_{STR}$ neurons in the unisensory trials at different locomotion states.

**d**, Neural trajectories of the $AC_{STR}$ neurons in the top three PCs of the stimulus subspace from the dPCA. Four colors in trajectories represent unisensory trial types. Solid lines, stationary conditions; dotted lines, moving conditions.

**e**,**f**, Projections distances of the unisensory trajectories (**e**) and distances between Go and No-go trajectories (**f**) at different locomotion states.

**g**, Mean classifier accuracies of auditory trials during 0.5 ~ 1 s post-stimulus period. S, stationary condition; M, moving condition.

**h**-**j**, Same as **c**,**e**,**f**, but for incongruent trials.

Data are presented as means ± SEM. Overlaid gray lines indicate individual data. For box plots, central lines indicate medians; boxes represent the $25^{th}$-$75^{th}$ percentiles; whiskers show maximum/minimum values excluding outliers. Source data are available in Source Data File. Sample numbers and statistical information are listed in Supplementary Data 1. NS, not significant; ***$P < 0.001$.
